# Metabolic Profiling of Aortic Stenosis and Hypertrophic Cardiomyopathy Identifies Mechanistic Contrasts in Substrate Utilisation

**DOI:** 10.1101/715680

**Authors:** Nikhil Pal, Animesh Acharjee, Zsuzsanna Ament, Tim Dent, Arash Yavari, Masliza Mahmod, Rina Ariga, James West, Violetta Steeples, Mark Cassar, Neil J. Howell, Helen Lockstone, Kate Elliott, Parisa Yavari, William Briggs, Michael Frenneaux, Bernard Prendergast, Jeremy S Dwight, Rajesh Kharbanda, Hugh Watkins, Houman Ashrafian, Julian L Griffin

## Abstract

**Background:** Aortic stenosis (AS) and hypertrophic cardiomyopathy (HCM) are highly distinct disorders leading to left ventricular hypertrophy (LVH), but whether cardiac metabolism substantially differs between these in humans remains to be elucidated.

**Method:** We undertook a detailed invasive (aortic root and coronary sinus) metabolic profiling in patients with severe AS and HCM in comparison to non-LVH controls, to investigate cardiac fuel selection and metabolic remodelling. These patients were assessed under different physiological states (at rest and during stress induced by pacing). The identified changes in the metabolome were further validated by metabolomic and orthogonal transcriptomic analysis, in separately recruited patient cohorts.

**Results:** We identified a highly discriminant metabolomic signature in severe AS characterised by striking accumulation of long-chain acylcarnitines, intermediates of long-chain transport and fatty acid metabolism, and validated this in a separate cohort. Mechanistically, we identify a down-regulation in the PPAR-α transcriptional network, including expression of genes regulating FAO.

**Conclusions:** We present a comprehensive analysis of changes in the metabolic pathways (transcriptome to metabolome) in severe AS, and its comparison to HCM. Our results demonstrate fundamental distinctions in substrate preference between AS and HCM, highlighting insufficient long-chain FAO, and the PPAR-α signalling network as a specific metabolic therapeutic target in AS.

## INTRODUCTION

As a continually working aerobic biological pump, the adult human heart has the highest energy demand for ATP per gram of tissue of any organ, requiring 6-30 kg daily(1). The healthy heart primarily relies on fatty acid β-oxidation (FAO), utilising circulating free fatty acids (FFA) or lipoprotein-derived triacylglycerols (50-70% ATP requirements), but also consumes carbohydrates (glucose/lactate/pyruvate), branched-chain amino acids and ketones(2). This metabolic flexibility enables the heart to meet physiological workloads.

The balance between cellular energy metabolism and contractile performance is disrupted in cardiac disease. Individuals with advanced chronic heart failure (HF) consistently display reduced cardiac high-energy phosphates (with up to 30% lower absolute cardiac [ATP])(3), and this is reproduced in animal HF models(4). Myocardial phosphocreatine:ATP ratio, an index of cardiac bioenergetic state, correlates with HF severity and strongly predicts mortality(5). Such observations highlight the energy-deplete state of the failing heart(6) and perturbations in cardiac intermediary energy metabolism.

Studies of the failing myocardium indicate substantial metabolic reconfiguration including: substrate utilisation switch from FFA to glucose(7), uncoupling of glycolysis from oxidation(8), FAO downregulation(9), impaired mitochondrial respiration(10) and decreased mitochondrial/cytosolic creatine kinase flux(11), together with loss of metabolic flexibility to stress(12). These are partly driven by nuclear receptor and transcriptional co-regulator signalling circuits orchestrating fuel selection and mitochondrial oxidative capacity, including: peroxisome proliferator-activated receptor α (PPARα)(13), a fatty-acid ligand binding master transcription factor promoting FAO; along with interacting regulators of oxidative metabolism retinoid X receptor α (RXRα)(14), estrogen-related receptor α (ERRα) and PPARy coactivator-la (PGC-1α)(15). Such changes have been conceptualised as a return to a fetal metabolic programme adapted to hypoxia, with preferential carbohydrate use and associated improvement in myocardial oxygen efficiency(16), but whether these become maladaptive with HF progression is unclear. Supporting a *causal* role for cardiac metabolic dysregulation, impaired FAO precedes systolic impairment in experimental models of left ventricular (LV) pressure overload(17); conversely, relief of haemodynamic stress restores oxidative metabolism before functional and structural recovery(18). Accordingly, therapies manipulating cardiac substrate utilisation have been trialled, including perhexiline(19), etomoxir(20) and trimetazidine(21), with variable impact on energetics and/or symptoms, through blockade of carnitine palmitoyltransferase 1 (CPT-1), which catalyses the rate-limiting step in mitochondrial β-oxidation, or β-oxidation enzymes.

In parallel with metabolic reconfiguration, a hallmark of the cardiac response to stress is structural remodelling, specifically left ventricular hypertrophy (LVH)(22). LVH can be primary in origin, exemplified by familial hypertrophic cardiomyopathy (HCM), or secondary to excessive haemodynamic loading (LV volume or pressure overload). Pressure overload-induced LVH, commonly resulting from aortic valvular stenosis (AS) or hypertension, is traditionally regarded as a compensatory mechanism normalising systolic wall stress(23) which, if persistent paradigmatically transitions to frank HF. LVH is associated with reduced myocardial high energy phosphate content(24) and consistently predicts adverse clinical outcomes including mortality and HF(25, 26). Inappropriately high LV mass is associated with higher cardiovascular risk in both HCM(27) and AS(28), while LVH regression through haemodynamic unloading (e.g. antihypertensive therapy) lowers cardiovascular risk(29). Morphologically, distinct LVH geometries have been identified which relate to cardiovascular risk(30), but without parallel progress in defining specific metabolic endotypes, a prerequisite for precision therapeutics targeting metabolism. We reasoned that analysis of the metabolome – reflecting a convergence of multiple levels of biological organisation and being functionally proximate to disease – in distinct LVH aetiologies could address this shortfall to: (i) identify discriminant metabolic endotypes; (ii) provide disease-specific mechanistic insights; and (iii) generate hypotheses for translational testing(31).

## RESULTS

### A targeted in vivo metabolomics strategy to probe human myocardial metabolism at baseline and during increased workload

To obtain a systematic quantitative assessment of cardiac metabolism *in vivo* in AS versus HCM populations, we undertook targeted plasma metabolic profiling using liquid chromatography tandem mass spectrometry (LC-MS/MS) to provide a comprehensive snapshot of substrate selection and metabolism (**Fig. 1a**). These patient groups were compared to a group that had no valvular heart disease or cardiomyopathy, and following an angiogram for suspected coronary artery disease, which turned out to be negative, were defined as a group free of macrovascular cardiovascular disease. These were referred to as the control group although microvascular ischaemia could not be excluded (32, 33). We focused on FAO, inferred from levels of acylcarnitine species, as well as intermediates from glycolysis, TCA cycle, amino acid, ketone and 1-carbon metabolism. To analyse myocardial substrate utilisation, we evaluated arteriovenous (AV) metabolite extraction by paired sampling of blood from the aortic root (AR, reflecting myocardial arterial blood supply) and coronary sinus (CS, representing cardiac venous effluent). Simultaneously samples from femoral venous (FV) blood to assess the ability of the peripheral metabolome to reflect myocardial biochemical alterations, as described for some metabolites(34). We further explored the ability of selectively increasing myocardial work to impact on the metabolomic phenotype and refine metabolomic signature by repeating sampling after right ventricular pacing.

**Figure 1.**
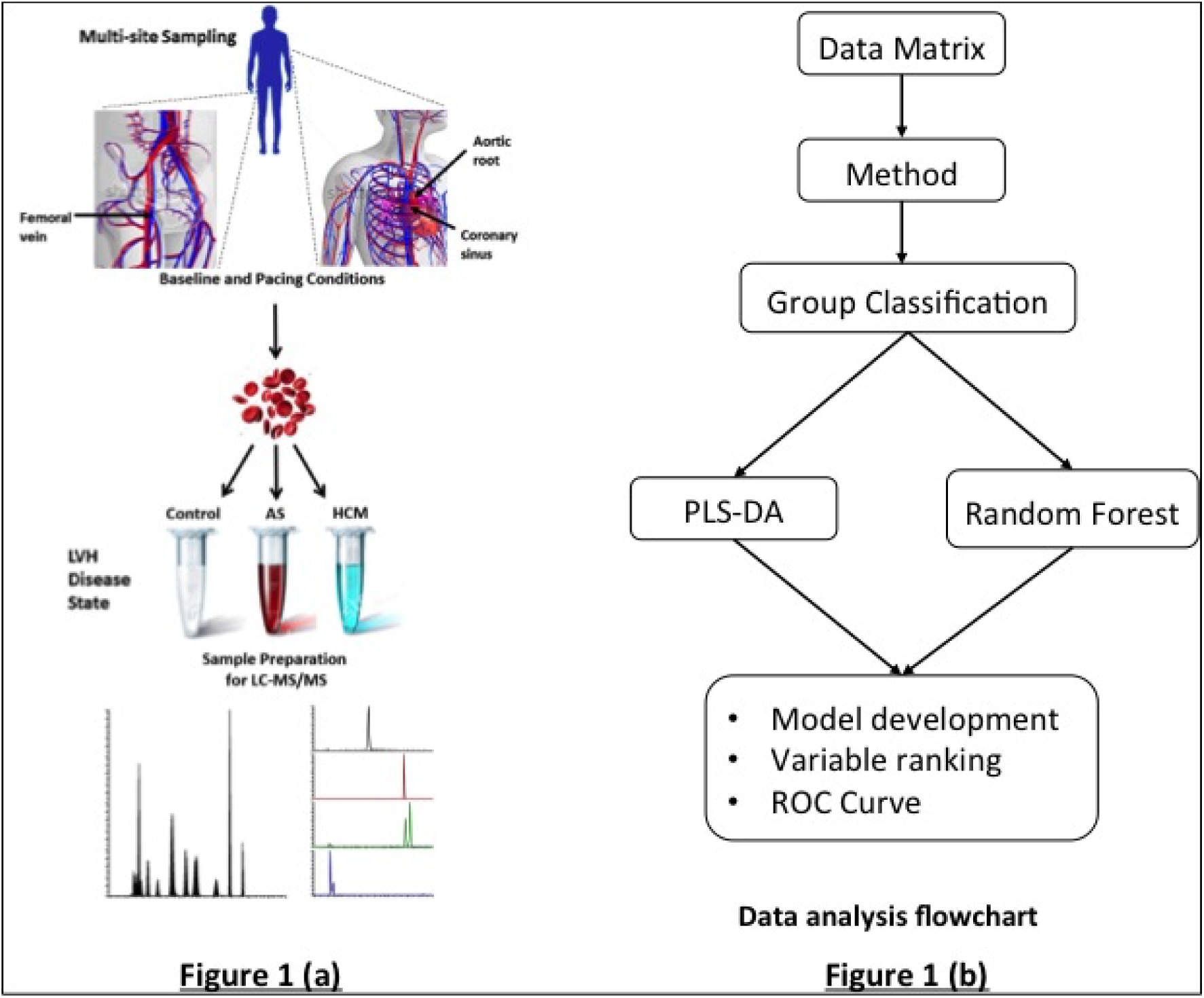
Illustration of the workflow for the study, **(a)** This details the collection of blood samples from multiple sites in the human body and processing using mass spectrometry platforms **(b)** Overview of multivariate statistics used in the analysis of the data.

This protocol was applied to a discovery cohort of AS (n = 36), HCM (n = 35) and control (n = 38, individuals undergoing elective cardiac catheterisation for suspected coronary artery disease) patients. Reflecting its natural history, AS subjects were older than HCM or control groups (mean 80 ± 7 years versus 62 ± 10 or 63 ± 10, respectively), and had higher incidence of hypertension and thus antihypertensive use (**Table 1** and **Supplementary Tables 1 and 2**). Notably, on transthoracic echocardiography, all three groups showed evidence of left ventricular hypertrophy, although the extent was substantially less in the control group than in either the AS or HCM groups.

**Table 1.** Table details the clinical and demographic characteristics of the controls, aortic stenosis (AS) and hypertrophic cardiomyopathy (HCM) cohorts. Differences between the cohorts were examined statistically using ANOVA with Bonferroni’s post-test for multiple comparisons. Statistical significance was defined as a *P <* 0.05.

|  | Controls<br>(n = 38) | Aortic stenosis<br>(n = 36) | HCM<br>(n = 35) | P value |
| --- | --- | --- | --- | --- |
| Age (years) | 63 ± 10 | 80 ± 7 | 62 ± 10 | <0.0001 |
| Men, n % | 79 | 53 | 71 | 0.04 |
| BMI (kg/m<sup>2</sup>) | 26.1 (23.7 - 28.5) | 28.1 (27.2 - 30.8) | 27.5 (25.9 - 31.7) | 0.06 |
| NYHA I | 14 (36%) | 3 (8%) | 6 (17%) | 0.008 |
| NYHA II | 18 (47%) | 19 (53%) | 26 (74%) | 0.05 |
| NYHA III | 6 (16%) | 11 (31%) | 3 (9%) | 0.05 |
| NYHA IV | 0 | 3 (8%) | 0 |  |
| PMH, n % |  |  |  |  |
| Hypertension | 50 | 75 | 40 | 0.009 |
| Diabetes | 16 | 25 | 11 | 0.3 |
| Dyslipidemia | 53 | 75 | 54 | 0.09 |
| COPD / Asthma | 13 | 14 | 14 | 0.9 |
| Family History | 61 | 44 | 69 | 0.1 |
| Medications, n % |  |  |  |  |
| Aspirin | 68 | 42 | 57 | 0.07 |
| Clopidogrel | 16 | 3 | 17 | 0.1 |
| ACE-I/ARB-II | 47 | 75 | 14 | <0.0001 |
| Beta-blockers | 40 | 47 | 49 | 0.7 |
| Statin | 66 | 61 | 46 | 0.2 |
| Calcium blockers | 24 | 28 | 40 | 0.3 |
| Diuretics | 16 | 72 | 11 | <0.0001 |
| OHA | 8 | 19 | 9 | 0.2 |
| Insulin | 11 | 3 | 0 | 0.08 |

*Values are mean ± SD, percentages or median (quartiles 1 to 3). BMI = Body mass index; PMH = past medical history; ACE = angiotensin-converting enzyme-inhibitors; IHD = ischemic heart disease; ARB = angiotensin-receptor antagonist-II. OHA = Oral hypoglycaemic agent.*

### Trans-myocardial arteriovenous extraction differences in LVH versus control

The trans-myocardial AV gradient of selected key carbon substrates (glucose, lactate, glutamate and alanine) was first to be measured and the results from this were consistent with their previously reported role in cardiac metabolism at rest and during stress (**Supplementary Fig. 1**)(35, 36). We next extended our analysis of trans-myocardial AV gradient to the complete metabolite profile for the cohort at baseline (**Supplementary Table 3**) and during pacing (**Supplementary Table 4**). This revealed significant cardiac extraction of glutamate, α-ketoisovalerate, keto-isoleucine, keto-leucine and pyruvate, but elution of succinate, in controls and AS at baseline. This pattern was largely recapitulated with pacing, with additional extraction of β-aminoisobutyric acid (BAIBA)(37) in AS. In the HCM cohort we found net extraction of glutamate and elution of succinate, regardless of pacing. Notably, compared to controls, both LVH cohorts displayed reduced (by ∼2/3) AV extraction of glutamate - a key anaplerotic substrate contributing to cellular oxidative capacity.

### Distinct metabolic profiles discriminate between AS and HCM

The differences between cohorts across the three collection sites were next analysed (**Figure 1b**). Models were built to determine the most discriminating metabolites using individual collection sites separately and then in combination. An initial PLS-DA comparison of the three groups using the combined baseline and pacing samples from all three-collection sites generated a robust discriminating model, suggesting that disease status had more influence on the dataset rather than site of collection or pacing status (**Fig. 2a,b**). An improved PLS-DA model was generated by comparing AS versus HCM and control combined as a single group (**Fig. 2c,d**). This separation was driven by increases in C14-OH and C18-OH acylcarnitines, and aconitate, and a decrease in α-ketoisocaproate, in the AS cohort. Similar results were obtained whether using metabolite levels from baseline or pacing samples separately or in combination; accordingly, further comparisons were made using the baseline samples only.

**Figure 2.**
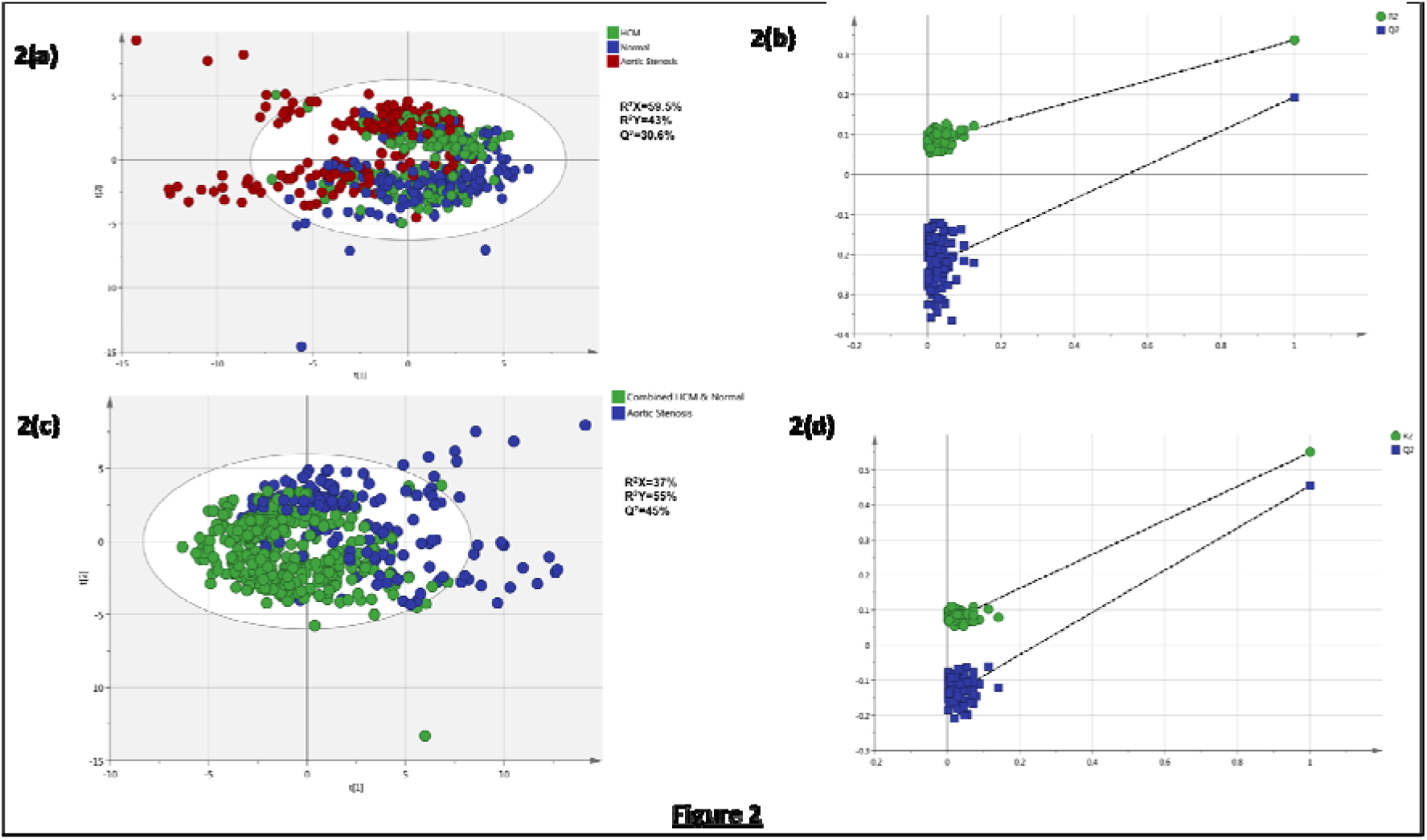
Score and validation plots showing comparison of aortic stenosis, HCM and control cohorts. **(a)** Score plot comparing all three cohorts. A robust model was built discriminating all three cohorts using PLS-DA (R^2^X=59.5%, R^2^Y=43%, Q^2^=30.6%). **(b)** Validation plot comparing all three cohorts a robust model was built discriminating all three cohorts using PLS-DA. **(c)** Score plot comparing the aortic stenosis cohort with the combined samples from HCM and control cohorts. A robust model was built using PLS-DA (R^2^X=37%, R^2^Y=55%, Q^2^=45%). **(d)** Validation plot comparing the aortic stenosis cohort with the combined samples from the HCM and control cohorts. The main discrimination was between aortic stenosis samples and the other two cohorts combined together.

Next, a pair-wise comparison of the three groups using combined samples from all three sites was performed. This generated robust models discriminating: HCM versus control (**Fig. 3a**); AS versus control (**Fig. 3b**); and AS versus HCM (**Fig. 3c**). All models passed the random permutation analysis and model misclassification, performed to assess their validity and goodness of fit (**Fig. 3d** and **Supplementary Fig. 2**). AS was clearly discriminated from both HCM and control cohorts: 94% correct across 362 samples and 90% correct across 370 samples, respectively.

**Figure 3.**
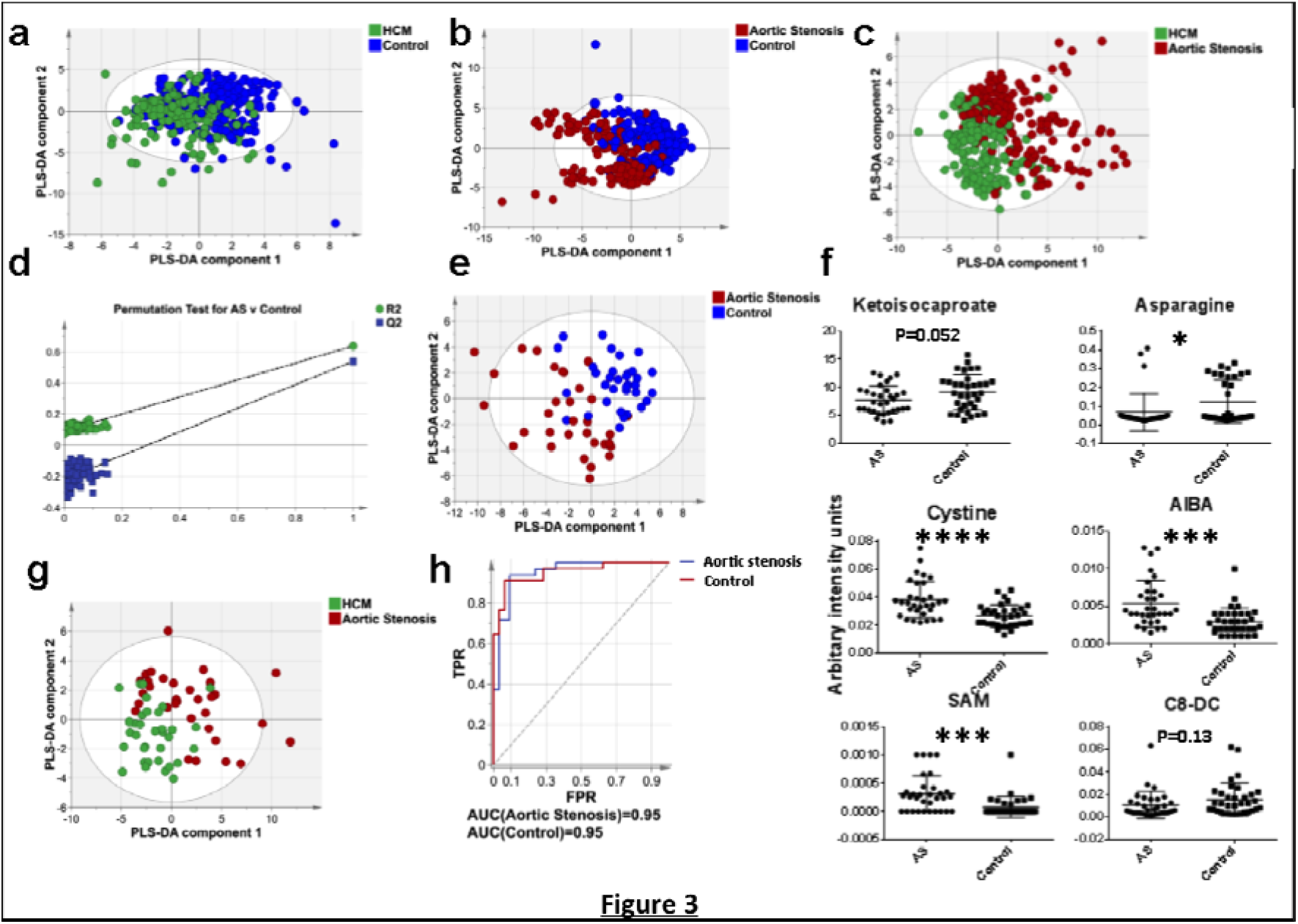
Multivariate pairwise-analysis of patient groups. **(a)** PLS-DA comparison between the HCM and control groups using blood samples from all three sampling sites (CS, AR, FV) (R^2^X=36%, R^2^Y=41%, Q^2^=18%). **(b)** PLS-DA comparison between the AS and control groups using blood samples from all three sampling sites (CS, AR, FV) (R^2^X=38%, R^2^Y=64%, Q^2^=54%). **(c)** PLS-DA comparison between the HCM and AS groups using blood samples from all three sampling sites (CS, AR, FV) (R^2^X=44%, R^2^Y=70%, Q^2^=53%) **(d)** Random permutation test of the model validity for the model in **b. (e)** PLS-DA comparison between the control and AS groups using blood samples from the CS (R^2^X=30%, R^2^Y=59%, Q^2^=37%). **(f)** Box-whisker plots of key metabolites that are discriminatory between the AS and control groups for the model in **e. (g)** PLS-DA comparison between the HCM and AS groups using blood samples from the CS (R^2^X=29%, R^2^Y=47%; Q^2^=18%). **(h)** ROC analysis of the predictive capability of the model in **e** discriminating between AS and control groups.

Variable Importance in Projection (VIP) scores were then used to rank the relative discriminatory value of metabolites responsible for the classification of AS versus control (37), HCM versus control (31) and AS versus HCM (41) comparisons (**Supplementary Fig. 3**). Of these the following metabolites were common to all three comparisons: isocitrate, proline, valine, carnitine, C2, C10, C14:1, C14-OH, C16, C16-OH and C18-OH, while some metabolites were disease-specific: AS versus control (asparagine, carnitine, ornithine and, C6-DC and C6 acylcarnitines) and HCM versus control (alanine, betaine, leucine, methionine and, C18:2-OH, C16:1-OH, and C16:2 acylcarnitines). PLS-DA analysis examining pairwise comparisons from CS, FV and AR samples separately was then performed. The most discriminating models were built using the CS samples, where robust models discriminating AS from control (**Figure 3e**) and AS from HCM (**Figure 3g**) were built. The model discriminating AS from control passed the random permutation test, CV-ANOVA (*P =* 1.2 × 10^−5^) and produced receiver operator control (ROC) curve with AUC = 0.95 (**Figure 3h**). This classification was driven in part by increase in cystine, BAIBA and S-adenosyl methionine, and decreases in C8-DC acylcarnitine, asparagine and α-ketoisocaproate in the AS cohort (**Fig. 3f**). The discrimination between AS and HCM cohorts was driven by greater concentrations of malate and, C14-OH, C20:1 and C12 acylcarnitines, and lower concentration of α-ketoisocaproate, uridine and histidine in the AS cohort (data not shown). Similar analyses were also performed on the FV and AR samples (**Supplementary Fig. 4-7**), which produced similar, albeit less robust models as compared to the CS. The significant age difference between subjects in the AS and the other cohorts was investigated by stratifying for age, to investigate whether this had an impact on the multivariate models constructed. Subjects over the age of 70 years were selected, and samples from all three collection sites collected both at baseline and pacing were included (**Supplementary Figure 8a**). A robust model discriminating AS and control cohorts was generated (**Supplementary Figure 8b**), which was validated using random permutation analysis (**Supplementary Figure 8c**). Also, several of the most discriminating metabolites were common between models built pre- and post-age stratification (**Supplementary Figure 8d**). Similarly, models discriminating control versus HCM and AS versus HCM cohorts for subjects over 70 years of age were also assembled.

To further validate the performance of the pairwise classification models, we used the selected metabolites to build random forest (RF) models and generated ROC curves (**Supplementary Fig. 9**). This analysis demonstrated good predictive ability of the selected metabolites to differentiate AS from the control cohort (**Fig. 4a-c**), associated with accumulation of long chain acylcarnitines in the AS cohort, suggesting a downregulation of FAO. On a three-way analysis between cohorts, the long chain acylcarnitines C16:1 and C18-OH were present in incrementally higher concentrations (significantly different between the pairs) between the control, HCM and AS cohorts, in the FV and AR collection sites (controls < HCM < AS).

**Figure 4.**
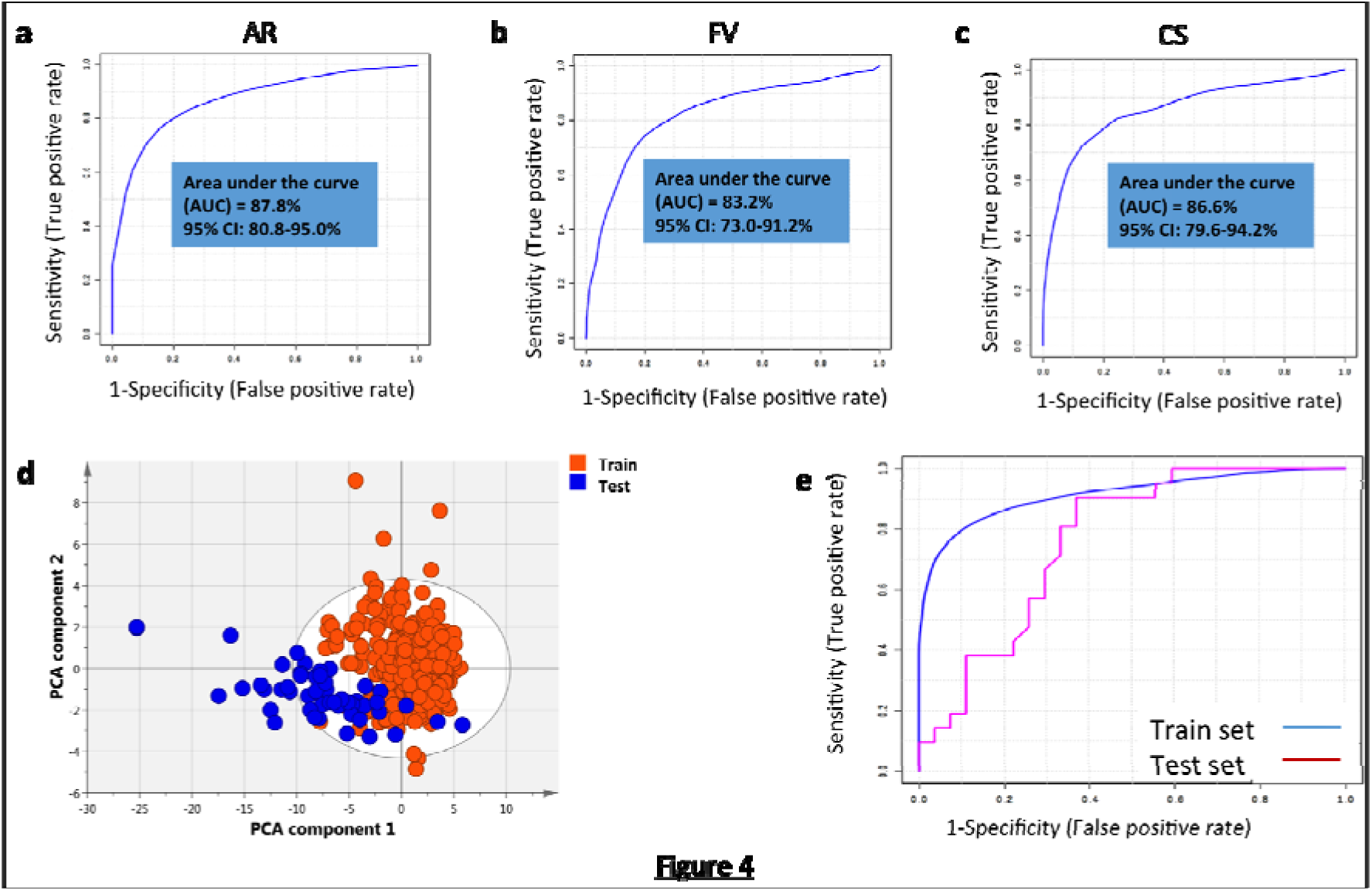
Multivariate and ROC curve analysis comparing the discovery and validation cohorts. **(a-c)**: ROC curve analysis comparing control and aortic stenosis cohorts from the discovery group. The analysis is performed individually for samples collected from the three sites: AR (Aortic root), FV (Femoral vein) and CS (Coronary sinus), **(d)** PCA scores plot of AS and control cohorts from the discovery and validation groups (for all metabolites combined), **(e)** ROC curves analysis comparing the FV samples from the discovery group (blue line) to the peripheral vein samples from the validation group (purple line).

Results from the discovery cohort comparing the AS and control cohorts were validated by analysing samples from the validation cohort. Compared to the discovery cohort, subjects with AS in the validation cohort were younger (71 vs 81 years, *P <* 0.0001), while controls in both cohorts were well matched for age (62 vs 61 years, *P =* 0.09). All subjects had normal LV ejection fraction and the majority were male (**Supplementary Table 5**). Exploratory principal components analysis (PCA) on all samples (AS and control combined) displayed clear separation between the discovery and validation cohorts (**Fig. 4d**). However, application of the multivariate model derived from the discovery cohort to the validation cohort discriminated AS and control cohorts as assessed by ROC analysis (AUC = 0.76, **Fig. 4e**).

### The cardiac transcriptomic profile of severe Aortic stenosis

To gain a more integrated understanding of the cellular consequences of AS-induced LVH and the mechanistic relevance of our plasma-based metabolomic findings, we used an orthogonal approach to examine the cardiac transcriptome in AS. We used whole-genome microarrays (Illumina) to profile intra-operative LV biopsies from 17 individuals with severe AS undergoing aortic valve replacement (AVR) in comparison to 13 control subjects (patients with coronary artery disease, normal LV systolic function and no significant aortic valve disease) undergoing coronary artery bypass grafting for ischaemic heart disease. The majority of patients in both groups were male, with no difference in age (71 vs 68 years, *P =* 0.84) (**Supplementary Table 6**).

Differentially expressed genes were identified using the Linear Models for Microarray Analysis (limma) package.(38) Of 19205 genes that passed filtering criteria, 8415 genes were significantly altered in AS at a false discovery rate (FDR) of 5%. Pathway enrichment analysis was performed using Ingenuity Pathway Analysis (IPA) on a subset of the differentially expressed genes (3656 had adjusted *P <* 0.001). Of 453 canonical pathways, 158 were significantly enriched (*P <* 0.05) for genes found altered in AS. These included some important metabolic pathways such as Adipogenesis (ranked 2, *P* = 1.6×10^−8^), PPAR signalling (ranked 48, *P =* 2.4×10^−3^) and PPARa/RXRa activation (ranked 51, *P =* 2.8×10^−3^) (**Supplementary Table 7** and **Supplementary Fig. 10**).

Consistent with the notion of energetic privation in LVH, we observed significant changes in the expression of genes encoding subunits of the cellular energy sensor AMP-activated protein kinase (AMPK) in AS including downregulation of protein kinase, AMP-activated, beta 1 non-catalytic subunit (*PRKAB1*) and protein kinase, AMP-activated, gamma 1 non-catalytic subunit (*PRKAG1*) and upregulation of protein kinase, AMP-activated, gamma 2 non-catalytic subunit (*PRKAG2*) and protein kinase, AMP-activated, alpha 1 catalytic subunit (*PRKAA1*) (**Supplementary Table 8**). Supporting the potential functional relevance of these findings, mice deficient in α2 AMPK develop more marked LVH and poorer contractile performance in response to pressure overload(39).

We also found differential expression in genes regulating different steps of FAO, including decreased expression of solute carrier family 27 (fatty acid transporter), member 5 (*SLC27A5*) and fatty acid binding protein 6 (*FABP6*) genes involved in the transport of FA across the sacrolemmal membrane, and increased expression of acyl-CoA synthetase long-chain family member 3 (*ACSL3*), which encodes proteins that convert long-chain FA into their acyl-CoA derivatives via thio-esterification. Further investigation of genes involved in peroxisomal β-oxidation revealed increased expression of ATP binding cassette subfamily D member 3 (*ABCD3*), acyl-CoA oxidase 1, palmitoyl (*ACOX1*), enoyl-CoA, hydratase/3-hydroxyacyl CoA dehydrogenase (*EHHADH*) and hydroxysteroid (17-beta) dehydrogenase 4 (*HSD17B4*) genes. This was thus indicative of increased FAO via the peroxisomal β-oxidation route.

There was a decrease in expression of genes from the acyl-CoA thioesterase family (*ACOT1, ACOT4, ACOT7* and *ACOT8*), encoding proteins that hydrolyse the CoA thioester of FA. These FFA can then be transported out of the peroxisomes into the mitochondria for further oxidation. There was increased expression of *CROT,* encoding a protein which catalyses the conversion of acyl-CoA to acylcarnitine, allowing transfer of the acylcarnitines from peroxisome to mitochondria. Reviewing genes involved in mitochondrial β-oxidation, we found decreased expression of genes involved in transport of acylcarnitines across the mitochondrial membrane (carnitine palmitoyltransferase 1A (*CPT1A*), carnitine palmitoyltransferase 2 (*CPT2) and* solute carrier family 25 (Carnitine/Acylcarnitine Translocase), member 20 (*SLC25A20).* There was also decreased expression of acyl-CoA dehydrogenase family member 8 (*ACAD8*) and hydroxysteroid (17-beta) dehydrogenase 10 (*HSD17B10*).

Several genes involved with microsomal FAO were also found significantly downregulated: cytochrome P450, family 4, subfamily B, polypeptide 1 (*CYP4B1*), cytochrome P450, family 27, subfamily A, polypeptide 1 (*CYP27A1*) and cytochrome P450, family 4, subfamily X, member 1 (*CYP4X1).* Thus, there was a possible reduction in the amount of FAO via the mitochondrial and microsomal routes.

We identified further evidence of remodelled myocardial lipid metabolism in AS, with downregulation in multiple genes encoding proteins regulating lipid biosynthesis including ELOVL fatty acid elongase 1 (*ELOVL1*) catalysing long chain fatty acid elongation at the endoplasmic reticulum; diacylglycerol O-acyltransferase 2 (*DGAT2*) that catalyses triacylglycerol synthesis; 1-acylglycerol-3-phosphate O-acyltransferase 5 (*AGPAT5*) and 1-acylglycerol-3-phosphate O-acyltransferase 7 (*AGPAT7*), both involved in phospholipid synthesis. Despite the reduction in multiple PPARα target genes, we observed a modest upregulation in the expression of *PPARA* itself in AS patients (Log2-fold change 0.43, *P =* 0.0009).

We also identified significant changes in genes encoding enzymes which regulate glycolysis, including upregulation of 6-phosphofructo-2-kinase/fructose-2,6-biphosphatase 2 (*PFKFB2*), and downregulation of glucose 6 phosphatase, catalytic, 3 (*G6PC3*), phosphoenolpyruvate carboxykinase 2 (*PCK2*), pyruvate dehydrogenase kinase, isozyme 1 & 2 (*PDK & PDK2).* Aconitase 2 (*ACO2*), the mitochondrial enzyme catalysing inter-conversion of citrate to isocitrate during the TCA cycle, was downregulated.

This analysis of the transcriptome reveals multiple changes in the myocardial energy sensing and generation circuitry of AS patients, in particular downregulation of multiple components of FA metabolism (**Fig. 5**).

**Figure 5.**
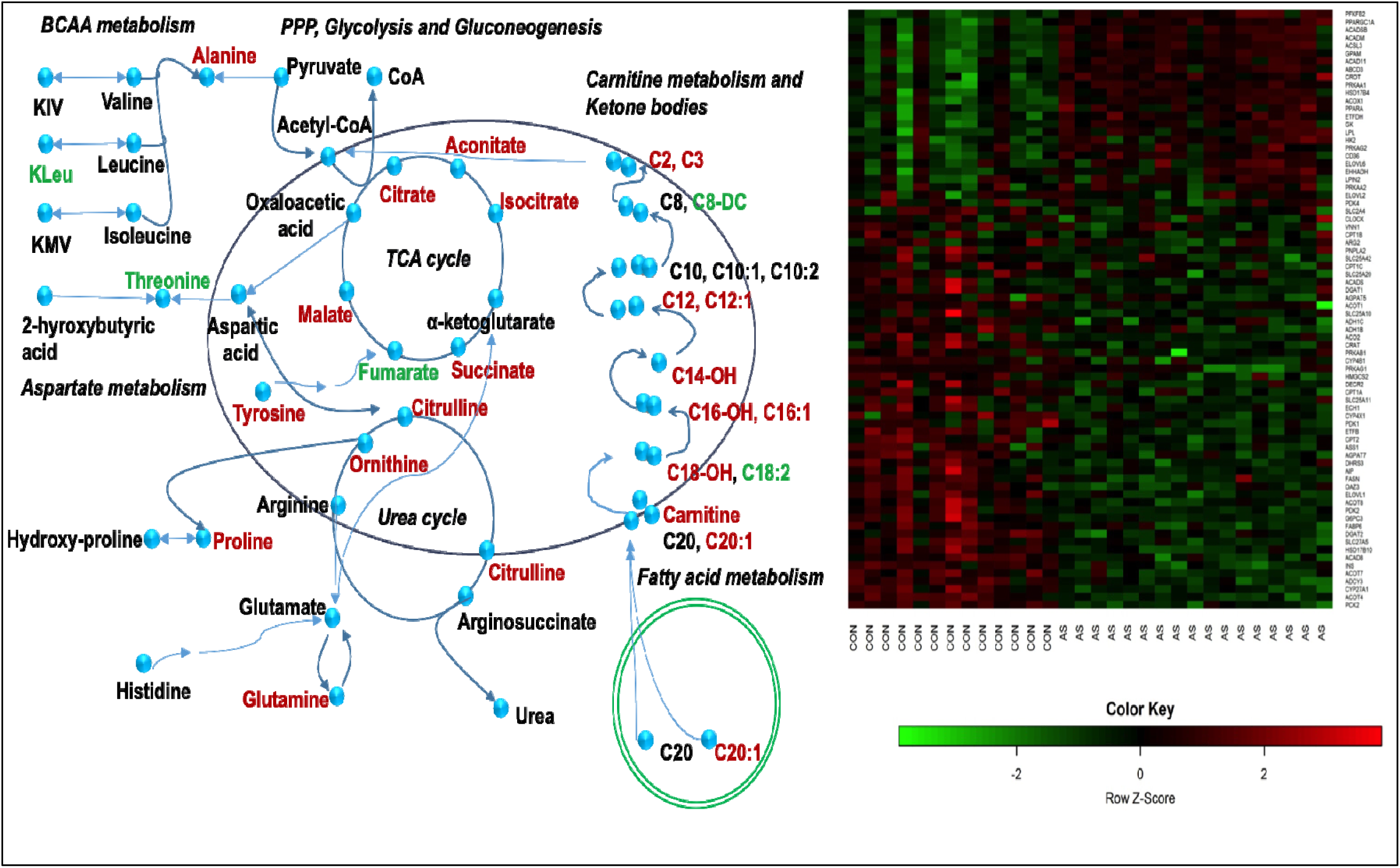
Overview map of impacted metabolic pathways and metabolite changes, along with the heat map of associated metabolism-related genes depicting their expression profiles, in aortic stenosis (AS) compared to controls. The concentration of the metabolite change was labelled green for a decreased concentration, red for increased concentration and black for no change in concentration. The heat map depicts expression patterns of 76 metabolism-related genes curated from the literature, of which 61 were significantly altered in AS patients, with the majority being down-regulated. The expression data are centred on zero for each gene with red indicating a sample has higher than average expression for a given gene while green indicates lower than average expression.

## DISCUSSION

Here, we leveraged a targeted metabolomic approach in well-defined human cohorts to draw biological insights into two common aetiologies of LVH: AS and HCM. Our systems medicine approaches disclosed a concordant bio-signature for reduced FAO in AS which, in the case of the plasma metabolome, was distinct from HCM.

Using a similar multi-site sampling approach to that recently described(40), we observe that predictive models to discriminate between AS, HCM and control populations performed best using samples obtained centrally (AR/CS) rather than peripherally (FV). However, the absolute reductions in AUCs were relatively small, enabling confirmation of the model from peripheral blood in a separate validation cohort.

By withdrawing paired blood samples from the AR and CS, we were able to directly examine trans-myocardial metabolite gradients *in vivo* in human(41). We found that the heart is a net importer of glutamate, α-ketoisovalerate, keto-isoleucine, keto-leucine and pyruvate, while it elutes succinate, malate and 3-hydroxybutyrate. The import of glutamate and metabolites of branch chain amino acids indicate the heart undergoes significant anaplerosis in order to maintain the citric acid cycle. Contrasting with recent studies highlighting a role for increased myocardial ketone body consumption as an alternative substrate to FAO in human and rodent failing hearts(42), we observed a net elution of β-hydroxybutyrate across control, AS and HCM groups. Bedi and colleagues(42) compared biopsies from end-stage human failing hearts at the time of orthotopic transplantation or LV assist device implantation with non-failing hearts from brain dead organ donors. They reported reductions in both cardiac lipid species and expression of enzymes involved in lipid metabolism including mid- and long-chain acylcarnitines, the high affinity carnitine transporter (*SLC22A5*)(42), but upregulation in β-hydroxybutyrate dehydrogenase 1 and 2 (*BDH1 and BDH2*) in end-stage HF, but not non-failing hearts (including those with LVH). Using mitochondrial proteomic profiling, Aubert and co-workers reported similar upregulation changes in murine models of both compensated LVH and HF, together with downregulation in enzymes involved in FAO, and a marked increase in BDH1 along with flux measurements consistent with increased oxidation of ketone bodies(43). Thus, the heart expresses the enzymes for producing ketone bodies. Combined with our findings, these data allude to specificity of metabolic rewiring in LVH versus advanced HF, and in AS there is a reduction in the oxidation of β-hydroxybutyrate that leads to its export from the heart. Somewhat surprisingly, as compared to individual collection sites the trans-myocardial gradients had a poor discriminatory power for disease state. Predictive models could be built examining samples from the CS, AR and FV that readily discriminated AS from the two other cohorts. This suggests that the classification models rely in part on systemic metabolism changes as much as those directly from the heart, suggesting AS should be understood as a systemic disease process(44).

Individuals with AS constituted the group most readily discriminated from normal subjects by plasma metabolomic profile. This was largely characterised by increases in long-chain acylcarnitine moieties, including C14, C16:1 and C18-OH, a metabolic signature suggesting incomplete FAO downstream of CPT-1, resonating with observations in advanced human HF(42). These long-chain acylcarnitine species are independently associated with maladaptive LV remodelling in patients with severe symptomatic AS(45). Similarly, our transcriptomic evaluation of LV biopsies from patients with AS highlighted the most prominent gene expression changes to affect lipid metabolism. Notably this included downregulation in specific components of mitochondrial β-oxidation, recently implicated as a likely contributor to the global acylcarnitine profile of severe HF(46). We further identified downregulation in cardiac-enriched subunit isoforms of the heterotrimeric master cellular energy gauge, AMPK, a key promoter of FA catabolism. AMPK’s actions include phosphorylation of the cytosolic enzyme acetyl-CoA carboxylase (ACC), reducing malonyl-CoA, thereby decreasing both the substrate for FA synthesis and simultaneously de-repressing CPT-1 activity to promote long-chain fatty acyl-CoA mitochondrial influx, in addition to influencing energy homeostasis and mitochondrial biogenesis via ERRα and PGC1-α(47).

Our finding of a large number of metabolic genes whose expression changes in AS, alludes to a significant cardiac metabolic reconfiguration in pressure overload-induced LVH, an observation mirrored in murine HF models(48) and potentially including altered mitochondrial protein acetylation, shown to involve FAO enzymes in HF(49). In agreement with our study, previous studies of HF have described increased long chain acyl carnitines in end-stage HF(50, 51) and pressure-overload induced HF(52), with these increases being reduced when patients are placed on circulatory support(51). Turer and co-workers employed paired-blood sampling to calculate that, following ischaemia-reperfusion, patients with left ventricular systolic dysfunction (LVSD) have reduced FFA uptake from the blood compared to controls(35). Yang and colleagues and Qiu and co-workers using the coronary artery ligation, MI induced-HF rat model, inferred alterations in FA biosynthesis and elongation from urinary metabolites and noted an increase in long-chain FAs in the blood, respectively(53, 54). Conversely, Lai and colleagues noted an increase in myocardial long-chain acyl-carnitine species and a decrease in L-carnitine in mice with HF(48). This may reflect differences in the staging of HF across patients and different model systems.

The current study has a number of potential limitations and caveats. Transcardiac metabolite gradient variability, especially with pacing, is not completely indicative of extent of associated changes in myocardial metabolism, because coronary sinus flow was not measured. The basis for accumulation of acylcarnitines in AS could represent, in part, saturation of capacity of CPT2, which functions to dissociate long chain fatty acids from carnitine on inner mitochondrial membranes. Finally, the “control” subjects used for obtaining myocardial biopsies were undergoing surgery for symptomatic coronary disease, and had mild LVH: thus this group cannot be entirely regarded as true “controls”.

In summary, we identify substantial differences in the plasma metabolic profiles accompanying two canonical forms of LVH, specifically that an acylcarnitine accumulation bio-signature indicative of repressed FAO associates strongly with human LV pressure overload represented by AS, supported by the finding of transcriptome reprogramming of FA metabolism. Our findings have important implications for attempts at metabolic modulation of LVH and the transition to HF, including in AS where, even in the setting of contemporary surgical or percutaneous valve replacement, myocardial remodelling(28, 55) and dysfunction(56) powerfully predict prognosis.

## METHODS

### Overall study design

#### Study Design

*We* undertook a prospective single centre observational study at the John Radcliffe Hospital, Oxford, UK to characterise the metabolome from 3 cohorts of patients: 1) HCM 2) AS with left ventricular hypertrophy and 3) controls. The study was granted ethical approval by National Research Ethics Service (NRES) Committee South Central–C (ethics reference number 13/SC/0155) and was monitored by the University of Oxford clinical trials and research governance department. Initial aim was to recruit an equal number (*n* = 35) of participants to each of the three cohorts, forming a total of 105 participants (discovery cohort). The results from the discovery cohort were then validated in a set of peripheral samples collected from a separate cohort of patients (validation cohort). These metabolomic findings were subsequently validated using an orthogonal technology, by performing transcriptomic analysis of the genes regulating these metabolic pathways, in LV biopsy samples from a separate cohort of AS and control patients.

#### Recruitment

Participants were aged 18 years or above with a known diagnosis of AS and HCM, and were undergoing angiography on the basis of prior determined clinical need. Controls were recruited as patients with clinically suspected coronary artery disease with normal systolic function, who were referred for diagnostic coronary angiography and in whom this revealed unobstructed coronary arteries. Patients then underwent baseline phenotyping, which included clinical history, examination, routine blood test, electrocardiogram (ECG) and transthoracic echocardiography (ECHO). Patients with severely impaired LV systolic function or conduction abnormality on ECG were excluded.

For transcriptomic analysis the LV biopsy samples were collected from patients with severe AS and controls (patients with ischaemic heart disease and normal LV systolic function) undergoing cardiac surgery.

#### Procedure

Following informed consent patients underwent cardiac catheterisation. Following coronary angiography plasma samples were collected at baseline and following right ventricular endocardial pacing from femoral vein (FV), coronary sinus (CS) and aortic root (AR). Pacing was commenced at 100 beats per minute for 60 sec and then incrementally increased every minute to a maximum of 140 beats per minute. Pacing was undertaken for a total duration of 3 minutes or stopped sooner if patient developed chest pain or ECG changes suggestive of ischaemia. In the validation cohort peripheral blood samples were obtained from a vein in the arm.

The blood samples were immediately placed on ice and then centrifuged at 3500 rpm at 4^°^C for 15 minutes. The resulting supernatant was separated and aliquoted into Eppendorf tubes and was re-centrifuged at 5000 rpm for 5 minutes. The supernatant was then aliquoted and stored at −80^°^C.

### Carnitine and aqueous metabolite analysis

HPLC grade solvents were obtained from Sigma Aldrich (Gillingham, Dorset, UK). All standards for optimisation and quantitation were obtained from Sigma Aldrich (Gillingham, Dorset, UK) with the exception of the internal standards, U-^13^C, ^15^N-glutamine, Di_10_-Leucine, D_8_-Phenylalanine, D_8_-Valine, U-^13^C, ^15^N-Proline which were acquired from Cambridge Isotope Laboratories (Andover, MA, USA).

#### Aqueous Metabolite Analysis

Samples were aliquoted (25 μL) to 96 well plates and proteins were precipitated using 400 μL MeOH containing a mixture of five internal standards at 2.5 μM. The precipitated samples were placed on a plate shaker for 5 min at 1000 rpm, and, using a liquid handling robot (Viaflo 96, Integra Biosciences, Nottingham, UK) transferred onto 96 well 0.66 mm glass fibre polystyrene filter plates (Corning Inc., NY, USA). Samples were filtered using a positive pressure 96 processor (Waters Ltd., Elstree, Hertfordshire, UK.) collected into a 1 mL polypropylene deep well plate (Thermo Scientific, Hemel Hempstead, Hertfordshire, UK.) and dried using an EZ-2 Evaporation system (Genevac Ltd., Ipswich, Suffolk, UK.). Samples were re-constituted in 100 μL of water containing 10 mM ammonium acetate, transferred to clean glass coated 200 μL 96-well plates, sealed using pre-slit foils and placed on the auto sampler. From each sample well 2 μL sample was injected.

Chromatographic analyses were performed using a Thermo Scientific Dionex UltiMate 3000 system (Thermo Scientific, Hemel Hempstead, Hertfordshire, UK), with an ACE C18-PFP 3 μm column (2.1 × 150 mm) (Advanced Chromatography Technologies Ltd., Aberdeen, UK) coupled to a TSQ Quantiva Triple Quadrupole Mass Spectrometer (Thermo Scientific, Hemel Hempstead, Hertfordshire, UK). The mobile phase gradient was run at 0.5 mL/min using water (mobile phase A) and acetonitrile (mobile phase B). The gradient started 100% A, and increased to 60% B from 1.6 to 4.5 minutes, followed by re-equilibration for 2 minutes. Data were acquired in both positive and negative ionisation modes using a capillary spray voltages of 3.5 kV and 2.5 kV, respectively. The ion transfer tube was set to operate at 350 °C whilst the vaporizer temperature was set to 400 °C. Sheath, auxiliary and sweep gases were set to 50, 15 and 2 arbitrary units, respectively.

#### Acylcarnitine analysis

Acyl-carnitines were measured according to the method described by Roberts *et al.* (57). Briefly, 100 μL internal standard solution mix (1.63 μM D_9_-free carnitine, 0.3 μM D_3_-acetyl carnitine, 0.06 μM D_3_-propionyl-carnitine, 0.06 μM D_3_-butyryl-carnitine, 0.06 μM D_9_-isovarelyl-carnitine, 0.06 μM D_3_-octanoyl-carnitine, 0.06 μM D_9_-myristoyl-carnitine, and 0.12 μM D_3_-palmitoyl-carnitine, Cambridge Isotope Laboratories, Andover, MA, USA) was added to 40 μL of the organic fraction of the methanol: chloroform extraction and the resulting mixture were dried down under nitrogen and derivatised with 100 μL of 3 M butanolic-HCI (Sigma-Aldrich, Louis, Missouri, USA). Samples were evaporated under nitrogen, re-constituted and sonicated in 4:1 acetonitrile: 0.1% formic acid in water before transferring them to autosampler vials.

Samples were analysed using an AB Sciex 5500 QTRAP mass spectrometer (AB Sciex UK Limited, Warrington, Cheshire) coupled to an Acquity UPLC system. Mobile phase A consisted of 0.1 % formic acid in water, while mobile phase B was acetonitrile. Two microliters of each sample was injected onto a Synergi Polar RP phenyl ether column (100 mm × 2.1 mm, 2.5 μm; Phenomenex, Macclesfield, Cheshire, UK). The analytical gradient started at 30 % B, followed by a linear increase to 100 % B over 3 min. The gradient was then held at 100 % B for 5 min, after which it was returned to the re-equilibration level of 30 % B for 2 min. A flow rate of 0.5 mL/min was used throughout. Data were analysed using the Quantitation Wizard within Analyst™ version 1.6 by AB Sciex Ltd. (Warrington, Cheshire, UK) and normalised against wet tissue weight and to the intensity of the internal standard.

### Statistical analysis and variable selection of the metabolomic data

We used two multivariate methods in these analyses to both maximise separations and also identify which metabolites are most important for these discriminations: partial least squares discriminant analysis (PLS-DA) and Random Forest (RF). As one of the primary objectives was to select (variable selection) the most important metabolites from the metabolites measured, a two-stage process was used. PLS-DA is a linear projection method, and all metabolites are assumed to be combined in a linear manner to maximise discrimination. In PLS-DA, the Variable Importance in Projection (VIP) scores were used to estimate the importance of each variable in the projection used in a PLS model^27^. RF does not assume any linearity and uses the sum of piecewise functions, and is thus able to discover more complex dependencies, and produce a more discrete set of metabolites.

In RF, we used a backward elimination process to select the best metabolites which result in good accuracy in terms of discrimination^23^. RF constructs a predictive model for the response using all predictors (carnitine and aqueous metabolites) and quantifies the importance of each carnitine and aqueous metabolites, in explaining the mis-classification error. We used a variable selection procedure to select the optimum number of carnitine and aqueous metabolites and using those selected metabolites RFs were iteratively fitted, so that they yielded the smallest out-of-bag (OOB) error rates. We further compared multivariate receiver operator control (ROC) curves to study how the number of selected variables impact on ROC performance.

Multivariate data analysis using PCA and PLS was performed in Simca 14.0 (Umetrics, Umeå, Sweden). RF classification of the classes on carnitine and aqueous metabolites and selection of carnitines and aqueous metabolites was conducted using the “randomForest” and “varSelRF”(58) package of R statistical software (https://www.r-project.org/).

### Transcriptomic analysis

#### Methods

Gene expression arrays: Total RNA was extracted from snap-frozen and pulverized heart tissue using the RNeasy Mini Kit (Qiagen, UK) following the manufacturers recommendations. Subsequent steps of cDNA synthesis, hybridisation, quality control, data normalisation and analysis were undertaken by the Genomics and Bioinformatics Core Facilities at the Wellcome Trust Centre for Human Genetics, Oxford. Illumina BeadChip microarrays (human WG-6 V2) were used according to manufacturer’s protocols.

Microarray data analysis: Raw intensity data were imported into the R statistical software (http://www.R-project.org) for further processing and analysis using BioConductor packages (59). Raw signal intensities were background corrected prior to being transformed and normalised using the ‘vsn’ package(60). A range of quality control checks were made on the data and it was filtered to remove genes not detected above background levels (detection score < 0.95) in at least 3 samples, resulting in a final dataset of 19205 genes and 30 samples (17 AS patients and 13 Controls).

Statistical analysis to identify differential gene expression was performed with the Linear Models for Microarray Analysis (limma) package (61). Raw p-values were corrected for multiple testing using the false discovery rate (FDR) controlling procedure of Benjamini and Hochberg (62). At 5% FDR, 8415 genes showed significant changes (adjusted *P <* 0.05) in their expression level between experimental groups.

## Author contributions

NP, AA, ZA, HW, HA and JLG designed the main study to follow metabolism in aortic stenosis and hypertrophic cardiomyopathy and wrote the manuscript. ZA, JAW and WB performed the metabolomics analysis. AA and JLG performed multivariate statistics on the metabolomic data. HA, VS, HL, KE performed transcriptional analysis on cardiac tissue. All authors read and approved the final manuscript. NP, AA and ZA share the first author position to reflect the fact that they led the human intervention study (NP), the metabolomic analysis (ZA) and the bioinformatics (AA).

## Acknowledgements

The research was supported by grants MR/P011705/1, MC_UP_A090_1006 and MR/P01836X/1. JLG is supported by the Imperial Biomedical Research Centre, NIHR. HA HW, NP are supported by the University of Oxford Biomedical Research Centre, NIHR. The double-blind, randomised study of perhexiline in AS was partially funded by an unrestricted grant from Sigma Pharmaceuticals, Sydney, Australia.

## References

1. Ashrafian H, and Neubauer S. Metabolomic profiling of cardiac substrate utilization: fanning the flames of systems biology? Circulation. 2009;119(13):1700–2.

2. Lopaschuk GD, Ussher JR, Folmes CD, Jaswal JS, and Stanley WC. Myocardial fatty acid metabolism in health and disease. Physiological reviews. 2010;90(1):207–58.

3. Neubauer S, Horn M, Pabst T, Harre K, Stromer H, Bertsch G, et al. Cardiac high-energy phosphate metabolism in patients with aortic valve disease assessed by 31P-magnetic resonance spectroscopy. Journal of investigative medicine: the official publication of the American Federation for Clinical Research. 1997;45(8):453–62.

4. Shen W, Asai K, Uechi M, Mathier MA, Shannon RP, Vatner SF, et al. Progressive loss of myocardial ATP due to a loss of total purines during the development of heart failure in dogs: a compensatory role for the parallel loss of creatine. Circulation. 1999;100(20):2113–8.

5. Neubauer S, Horn M, Cramer M, Harre K, Newell JB, Peters W, et al. Myocardial phosphocreatine-to-ATP ratio is a predictor of mortality in patients with dilated cardiomyopathy. Circulation. 1997;96(7):2190–6.

6. Neubauer S. The failing heart—an engine out of fuel. The New England journal of medicine. 2007;356(11):1140–51.

7. Doenst T, Nguyen TD, and Abel ED. Cardiac metabolism in heart failure: implications beyond ATP production. Circulation research. 2013;113(6):709–24.

8. Davila-Roman V, Vedala G, Herrero P, de las Fuentes L, Rogers J, Kelly D, et al. Altered myocardial fatty acid and glucose metabolism in idiopathic dilated cardiomyopathy. Journal of the American College of Cardiology. 2002;40(2):271.

9. Osorio JC, Stanley WC, Linke A, Castellari M, Diep QN, Panchal AR, et al. Impaired myocardial fatty acid oxidation and reduced protein expression of retinoid X receptor-alpha in pacing-induced heart failure. Circulation. 2002;106(5):606–12.

10. Sharov VG, Todor AV, Silverman N, Goldstein S, and Sabbah HN. Abnormal mitochondrial respiration in failed human myocardium. Journal of molecular and cellular cardiology. 2000;32(12):2361–7.

11. Weiss RG, Gerstenblith G, and Bottomley PA. ATP flux through creatine kinase in the normal, stressed, and failing human heart. Proceedings of the National Academy of Sciences of the United States of America. 2005;102(3):808–13.

12. Neglia D, De Caterina A, Marraccini P, Natali A, Ciardetti M, Vecoli C, et al. Impaired myocardial metabolic reserve and substrate selection flexibility during stress in patients with idiopathic dilated cardiomyopathy. American journal of physiology Heart and circulatory physiology. 2007;293(6):H3270.

13. Barger PM, Brandt JM, Leone TC, Weinheimer CJ, and Kelly DP. Deactivation of peroxisome proliferator-activated receptor-alpha during cardiac hypertrophic growth. The Journal of clinical investigation. 2000;105(12):1723–30.

14. Oka S, Zhai P, Yamamoto T, Ikeda Y, Byun J, Hsu CP, et al. Peroxisome Proliferator Activated Receptor-alpha Association With Silent Information Regulator 1 Suppresses Cardiac Fatty Acid Metabolism in the Failing Heart. Circulation Heart failure. 2015;8(6): 1123–32.

15. Sihag S, Cresci S, Li AY, Sucharov CC, and Lehman JJ. PGC-1alpha and ERRalpha target gene downregulation is a signature of the failing human heart. Journal of molecular and cellular cardiology. 2009;46(2):201–12.

16. Korvald C, Elvenes OP, and Myrmel T. Myocardial substrate metabolism influences left ventricular energetics in vivo. Am J Physiol Heart Circ Physiol. 2000;278(4):H1345–51.

17. Zhang L, Jaswal JS, Ussher JR, Sankaralingam S, Wagg C, Zaugg M, et al. Cardiac insulin-resistance and decreased mitochondrial energy production precede the development of systolic heart failure after pressure-overload hypertrophy. Circulation Heart failure. 2013;6(5):1039–48.

18. Byrne NJ, Levasseur J, Sung MM, Masson G, Boisvenue J, Young ME, et al. Normalization of cardiac substrate utilization and left ventricular hypertrophy precede functional recovery in heart failure regression. Cardiovascular research. 2016;110(2):249–57.

19. Abozguia K, Elliott P, McKenna W, Phan TT, Nallur-Shivu G, Ahmed I, et al. Metabolic modulator perhexiline corrects energy deficiency and improves exercise capacity in symptomatic hypertrophic cardiomyopathy. Circulation. 2010;122(16):1562–9.

20. Schmidt-Schweda S, and Holubarsch C. First clinical trial with etomoxir in patients with chronic congestive heart failure. Clinical science (London, England: 1979). 2000;99(1):27–35.

21. Fragasso G, Perseghin G, De Cobelli F, Esposito A, Palloshi A, Lattuada G, et al. Effects of metabolic modulation by trimetazidine on left ventricular function and phosphocreatine/adenosine triphosphate ratio in patients with heart failure. European heart journal. 2006;27(8):942–8.

22. Heusch G, Libby P, Gersh B, Yellon D, Bohm M, Lopaschuk G, et al. Cardiovascular remodelling in coronary artery disease and heart failure. Lancet (London, England). 2014;383(9932):1933–43.

23. Grossman W, Jones D, and McLaurin LP. Wall stress and patterns of hypertrophy in the human left ventricle. The Journal of clinical investigation. 1975;56(1):56–64.

24. Zhang J, Merkle H, Hendrich K, Garwood M, From AH, Ugurbil K, et al. Bioenergetic abnormalities associated with severe left ventricular hypertrophy. The Journal of clinical investigation. 1993;92(2):993–1003.

25. Levy D, Garrison RJ, Savage DD, Kannel WB, and Castelli WP. Prognostic implications of echocardiographically determined left ventricular mass in the Framingham Heart Study. The New England journal of medicine. 1990;322(22):1561–6.

26. de Simone G, Gottdiener JS, Chinali M, and Maurer MS. Left ventricular mass predicts heart failure not related to previous myocardial infarction: the Cardiovascular Health Study. Eur Heart J. 2008;29(6):741–7.

27. Spirito P, Bellone P, Harris KM, Bernabo P, Bruzzi P, and Maron BJ. Magnitude of left ventricular hypertrophy and risk of sudden death in hypertrophic cardiomyopathy. The New England journal of medicine. 2000;342(24):1778–85.

28. Cioffi G, Faggiano P, Vizzardi E, Tarantini L, Cramariuc D, Gerdts E, et al. Prognostic effect of inappropriately high left ventricular mass in asymptomatic severe aortic stenosis. Heart (British Cardiac Society). 2011;97(4):301–7.

29. Devereux RB, Wachtell K, Gerdts E, Boman K, Nieminen MS, Papademetriou V, et al. Prognostic significance of left ventricular mass change during treatment of hypertension. Jama. 2004;292(19):2350–6.

30. Garg S, de Lemos JA, Ayers C, Khouri MG, Pandey A, Berry JD, et al. Association of a 4-Tiered Classification of LV Hypertrophy With Adverse CV Outcomes in the General Population. JACC Cardiovascular imaging. 2015;8 (9): 1034–41.

31. Griffin JL, Atherton H, Shockcor J, and Atzori L. Metabolomics as a tool for cardiac research. Nature reviews Cardiology. 2011;8(11):630–43.

32. Johnson BD, Shaw, L.J., Pepine, C.J., Reis, S.E., Kelsey, S.F., Sopko, G., Rogers, W.J., Mankad, S., Sharaf, B.L., Bittner, V., Bairey Merz, C.N. Persistent chest pain predicts cardiovascular events in women without obstructive coronary artery disease: results from the NIH-NHLBI-sponsored Women’s Ischaemia Syndrome Evaluation (WISE) study. European Heart J. 2006;27(12):1408–15.

33. Bugiardini R. Women, ‘non-specific’ chest pain, and normal or near-normal coronary angiograms are not synonymous with favourable outcome. European Heart J. 2006;27:1387.

34. Makrecka M, Kuka J, Volska K, Antone U, Sevostjanovs E, Cirule H, et al. Long-chain acylcarnitine content determines the pattern of energy metabolism in cardiac mitochondria. Molecular and cellular biochemistry. 2014;395(1-2):1–10.

35. Turer AT, et al. Metabolomic profiling reveals distinct patterns of myocardial substrate use in humans with coronary artery disease or left ventricular dysfunction during surgical ischemia/reperfusion. Circulation. 2009;119(13):1736–46.

36. Thomassen A, Bagger JP, Nielsen TT, and Henningsen P. Altered global myocardial substrate preference at rest and during pacing in coronary artery disease with stable angina pectoris. The American journal of cardiology. 1988;62(10 Pt 1):686–93.

37. Roberts LD, Bostrom P, O’Sullivan JF, Schinzel RT, Lewis GD, Dejam A, et al. beta-Aminoisobutyric acid induces browning of white fat and hepatic beta-oxidation and is inversely correlated with cardiometabolic risk factors. Cell metabolism. 2014;19(1):96–108.

38. Gentleman R, Carey V, and Huber W. Bioinformatics and Computational Biology Solutions Using R and Bioconductor. 2006.

39. Zhang P, Hu X, Xu X, Fassett J, Zhu G, Viollet B, et al. AMP activated protein kinase-alpha2 deficiency exacerbates pressure-overload-induced left ventricular hypertrophy and dysfunction in mice. Hypertension (Dallas, Tex : 1979). 2008;52(5):918–24.

40. Lewis GD, Wei, R., Liu, E., Yang, E., Shi, X., Martinovic, M., Farrell, L., Asnani, A., Cyrille, M., Ramanathan, A., Shaham, O., Berriz, G., Lowry, P.A., Palacios, I.F., Taşan, M., Roth, F.P., Min, J., Baumgartner, C., Keshishian, H., Addona, T., Mootha, V.K., Rosenzweig, A., Carr, S.A., Fifer, M.A., Sabatine, M.S., Gerszten, R.E. Metabolite profiling of blood from individuals undergoing planned myocardial infarction reveals early markers of myocardial injury. Journal of Clinical Investigation. Journal of Clinical Investigation. 2008;118:3503–12.

41. Taegtmeyer H, Young ME, Lopaschuk GD, Abel ED, Brunengraber H, Darley-Usmar V, et al. Assessing Cardiac Metabolism: A Scientific Statement From the American Heart Association. Circulation research. 2016;118(10):1659–701.

42. Bedi KC Jr SN, Brandimarto J, Aziz M, Mesaros C, Worth AJ, Wang LL, Javaheri A, Blair IA, Margulies KB, Rame JE. Evidence for Intramyocardial Disruption of Lipid Metabolism and Increased Myocardial Ketone Utilization in Advanced Human Heart Failure. Circulation. 2016;133(8):706–16.

43. Aubert G MO, Horton JL, Lai L, Vega RB, Leone TC, Koves T, Gardell SJ, Krüger M, Hoppel CL, Lewandowski ED, Crawford PA, Muoio DM, Kelly DP. The Failing Heart Relies on Ketone Bodies as a Fuel. Circulation. 2016;133(8):698–705.

44. Otto CM, and Prendergast B. Aortic-Valve Stenosis — From Patients at Risk to Severe Valve Obstruction. New England Journal of Medicine. 2014;371(8):744–56.

45. Elmariah S, Farrell LA, Furman D, and, et al. Association of acylcarnitines with left ventricular remodeling in patients with severe aortic stenosis undergoing transcatheter aortic valve replacement. JAMA Cardiology. 2018;3(3):242–6.

46. Ruiz M, Labarthe F, Fortier A, Bouchard B, Thompson Legault J, Bolduc V, et al. Circulating acylcarnitine profile in human heart failure: a surrogate of fatty acid metabolic dysregulation in mitochondria and beyond. Am J Physiol Heart Circ Physiol. 2017;313(4):H768–h81.

47. Hu X, Xu X, Lu Z, Zhang P, Fassett J, Zhang Y, et al. AMP activated protein kinase-alpha2 regulates expression of estrogen-related receptor-alpha, a metabolic transcription factor related to heart failure development. Hypertension (Dallas, Tex: 1979). 2011;58(4):696–703.

48. Lai L, et al. Energy metabolic reprogramming in the hypertrophied and early stage failing heart: a multisystems approach. Circulation Heart failure. 2014;7(6):1022–31.

49. Horton JL, Martin OJ, Lai L, Riley NM, Richards AL, Vega RB, et al. Mitochondrial protein hyperacetylation in the failing heart. JCI insight. 2016;2(1).

50. Cheng ML, et al. Metabolic disturbances identified in plasma are associated with outcomes in patients with heart failure: diagnostic and prognostic value of metabolomics. J Am Coll Cardiol. 2015;65(15):1509–20.

51. Ahmad T, et al. Prognostic Implications of Long-Chain Acylcarnitines in Heart Failure and Reversibility With Mechanical Circulatory Support. J Am Coll Cardiol. 2016;67(3):291–9.

52. Sansbury BE, et al. Metabolomic analysis of pressure-overloaded and infarcted mouse hearts. Circulation Heart failure. 2014;7(4):643–42.

53. Yang D, et al. Urinary Metabolomic Profiling Reveals the Effect of Shenfu Decoction on Chronic Heart Failure in Rats. Molecules. 2015;20(7): 11915–29.

54. Qiu Q, et al. Plasma metabonomics study on Chinese medicine syndrome evolution of heart failure rats caused by LAD ligation. BMC Complement Altern Med. 2014;14:232.

55. Duncan Al, Lowe BS, Garcia MJ, Xu M, Gillinov AM, Mihaljevic T, et al. Influence of concentric left ventricular remodeling on early mortality after aortic valve replacement. The Annals of thoracic surgery. 2008;85(6):2030–9.

56. Dahl JS, Videbaek L, Poulsen MK, Rudbaek TR, Pellikka PA, and Moller JE. Global strain in severe aortic valve stenosis: relation to clinical outcome after aortic valve replacement. Circulation Cardiovascular imaging. 2012;5(5):613–20.

57. Roberts LD, West JA, Vidal-Puig A, and Griffin JL. Methods for Performing Lipidomics in White Adipose Tissue. Methods in Enzymology. 2014;538:211–31.

58. Diaz-Uriarte R, and De Andres SA. Gene selection and classification of microarray data using random forest. BMC bioinformatics. 2006;7(1):3.

59. Gentleman RC, Carey VJ, Bates DM, Bolstad B, Dettling M, Dudoit S, et al. Bioconductor: open software development for computational biology and bioinformatics. Genome Biology. 2004;5(10).

60. Huber W, von Heydebreck A, Sultmann H, Poustka A, and Vingron M. Variance stabilization applied to microarray data calibration and to the quantification of differential expression. Bioinformatics (Oxford, England). 2002;18 Suppl 1:S96–104.

61. Gentleman RC, V.; Huber, W.; Irizarry, R.; Dudoit, S;. Bioinformatics and Computational Biology Solutions Using R and Bioconductor. Springer-Verlag, New York 2005.

62. Benjamini Y, and Hochberg Y. CONTROLLING THE FALSE DISCOVERY RATE - A PRACTICAL AND POWERFUL APPROACH TO MULTIPLE TESTING. Journal of the Royal Statistical Society Series B-Methodological. 1995;57(1):289–300.

