## Supplementary Data for "Metabolic Profiling of Aortic Stenosis and Hypertrophic Cardiomyopathy Identifies Mechanistic Contrasts in Substrate Utilisation"

**Supplementary Data: Metabolic Profiling of Aortic Stenosis and Hypertrophic Cardiomyopathy Identifies Mechanistic Contrasts for Patient Treatment**

Nikhil Pal,^1,2*^ Animesh Acharjee,^3,4,5*^ Zsuzsanna Ament,^3,4*^ Tim Dent,^1^ Arash Yavari,^1,2^ Masliza Mahmod,^1^ Rina Ariga,^1^ James West,^3,4^ Violetta Steeples,^6^ Mark Cassar,^1^ Neil J. Howell,^7^ Helen Lockstone,^6^ Kate Elliott,^6^ Parisa Yavari,^1^ William Briggs,^3^ Michael Frenneaux,^8^ Bernard Prendergast,^1^ Jeremy S Dwight,^1^ Rajesh Kharbanda,^1^ Hugh Watkins,^1^ Houman Ashrafian^1,2#^ and Julian L Griffin^3,4#^

^1^Division of Cardiovascular Medicine, and ^2^Department of Experimental Therapeutics, Radcliffe Department of Medicine, John Radcliffe Hospital, Oxford, UK

^3^ Department of Biochemistry and Cambridge Systems Biology Centre, and ^4^MRC-Human Nutrition Research Unit, University of Cambridge, UK

^5^ Institute of Cancer and Genomic Sciences, Centre for Computational Biology, University of Birmingham, B15 2TT, UK

^6^ Wellcome Trust Centre for Human Genetics (WTCHG), University of Oxford, Oxford OX3 7BN, UK W

^7^ Department of Cardiothoracic Surgery, University Hospital Birmingham, Edgbaston, Birmingham, UK

^8^ Norwich Medical School, University of East Anglia, Bob Champion Research and Educational Building, James Watson Road, Norwich Research Park, Norwich, NR4 7UQ

^13^ Norwich Medical School, University of East Anglia, Bob Champion Research and Educational Building, James Watson Road, Norwich Research Park, Norwich, NR4 7UQ

*Contributed equally	^#^ Contributed equally

***For correspondence:***

Professor Houman Ashrafian	Professor Julian L Griffin,

Experimental Therapeutics	Department of Biochemistry,

Radcliffe Department of Medicine	University of Cambridge,

University of Oxford	Tennis Court Road,

John Radcliffe Hospital	CB2 1GA, UK

Oxford, OX3 9DU, UK

**Supplementary**

**Supplementary Figure 1** Trans-cardiac gradient measurements of glucose, lactate, glutamate and alanine. Glucose (A & B), lactate (C & D), glutamate (E & F) and alanine (G & H) concentrations in the aortic root and coronary sinus as measured in all patient groups combined together. The trans-cardiac gradient of glucose and lactate was measured for the first consecutive 61 patients. At baseline, as expected there was a significant negative AV gradient (from AR to CS) in glucose (-0.25 μmol/L, *P =* 0.0004), lactate (-0.07 μmol/L, *P <* 0.0001) and glutamate (-1.03 area ratio, *P* < 0.0001), consistent with resting cardiac uptake of these metabolites, but without differential concentration of alanine. While the concentration of glucose in the coronary sinus increased during pacing, it still remained lower as compared to the aortic root; indicating a net lower extraction compared to baseline. CS lactate concentrations were nominally higher post-pacing, consistent with net elution, but did not reach statistical significance. The trans-cardiac gradient of glutamate and alanine was measured for a total of 63 patients. There was a significantly lower concentration of glutamate in the coronary sinus both during baseline (*P* < 0.0001) and pacing (*P* < 0.0001), implying a net extraction of glutamate by the heart. There was no significant difference in the concentration of alanine between the aortic root and coronary sinus, both during baseline and pacing states. The concentrations are shown as box plots, with the box representing quartiles 1 to 3 and the whiskers representing 10 to 90 percentile.

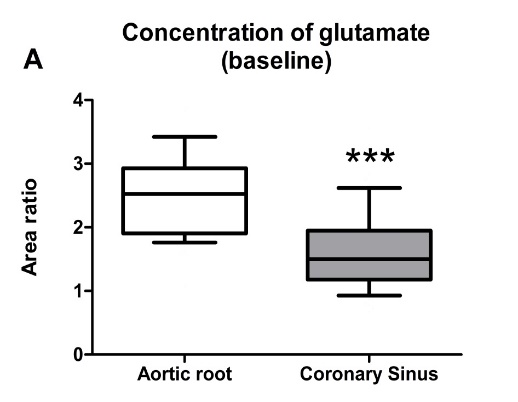

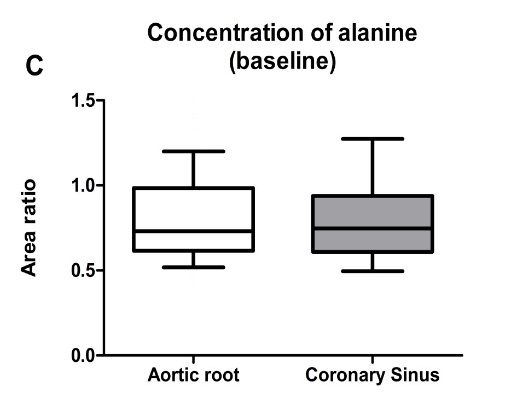

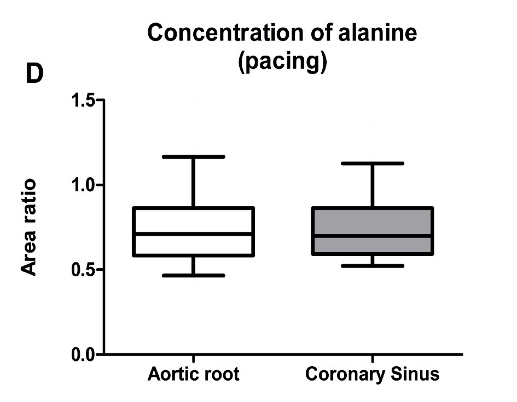

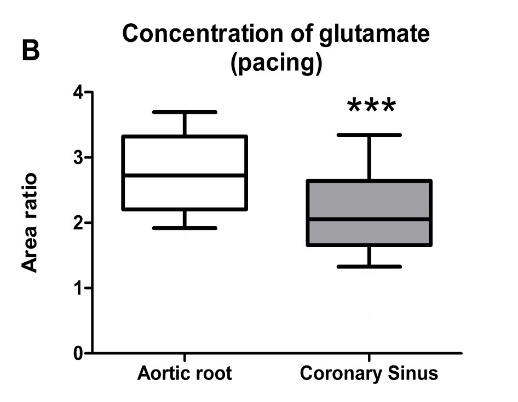

**E**

**F**

**H**

**G**

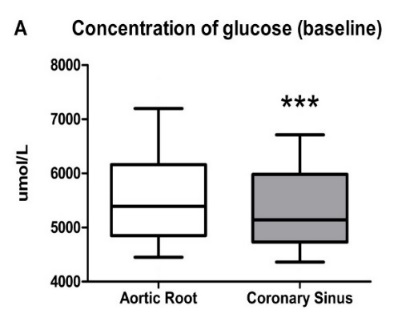

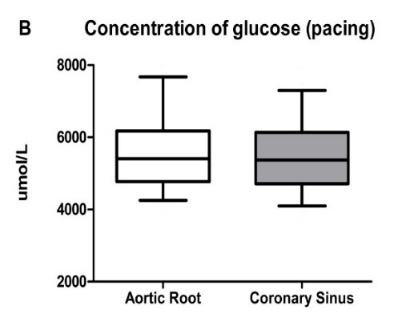

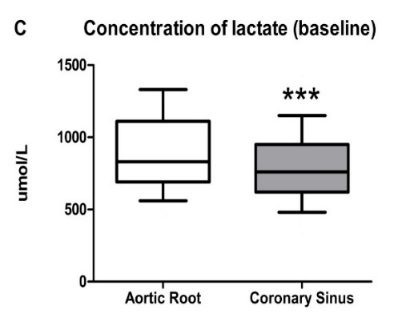

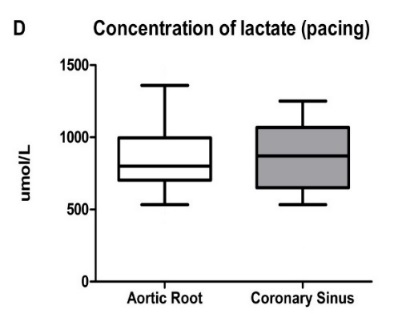

**Supplementary Figure 2** The corresponding misclassification tables for the models in Figure 2 are shown to display the number of samples in each group, and the percentage values of the correctly classified samples.

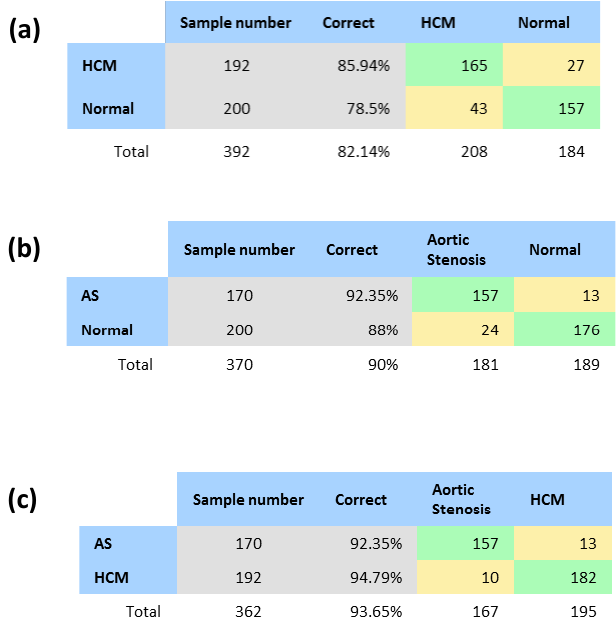

**Supplementary Figure 3** Further analysis of most discriminatory metabolites. (**a**) Venn diagram of the most discriminatory metabolites for the two group comparisons (both increased and decreased metabolites are shown for the comparisons). (**b**) Metabolites primarily responsible for the classifications using Variable Importance in Projection (VIP) scores to rank their relative discriminatory value (identifying metabolites with a VIP score of > 1 that were common and unique).

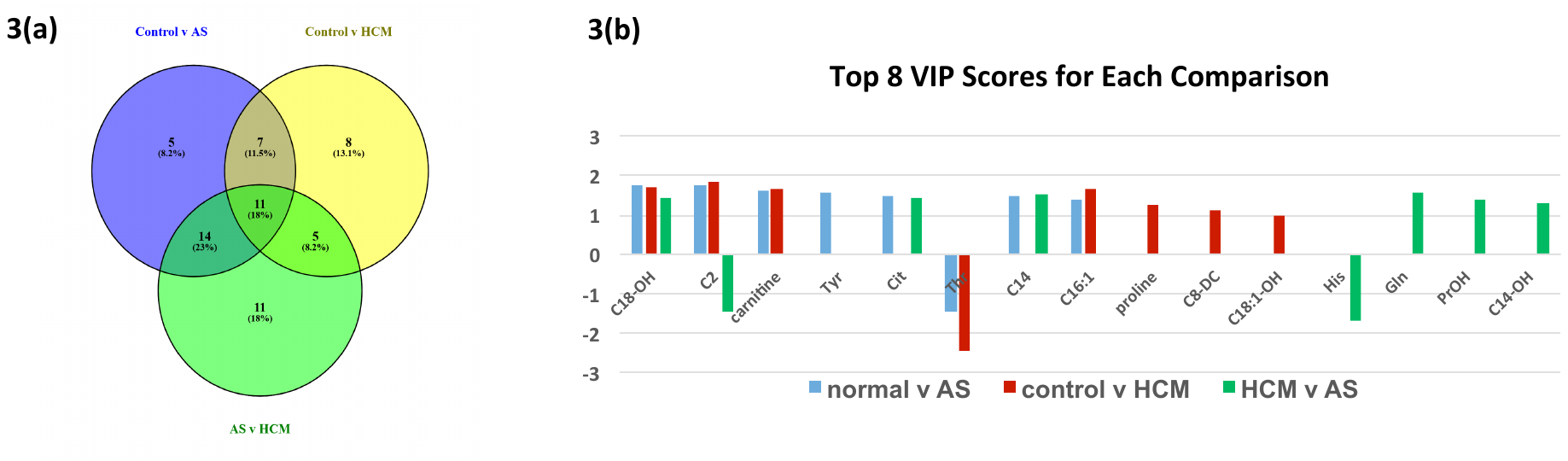

**Supplementary Figure 4 Metabolites** selected using random forest and their changes are shown in the box plot for control vs. aortic stenosis: a) AR site b) FV site c) CS site**.**

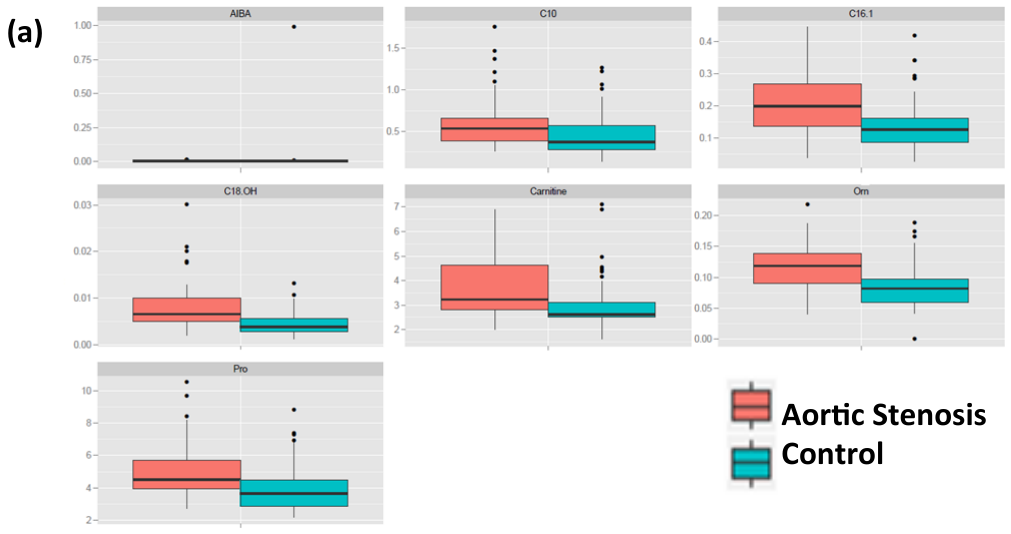

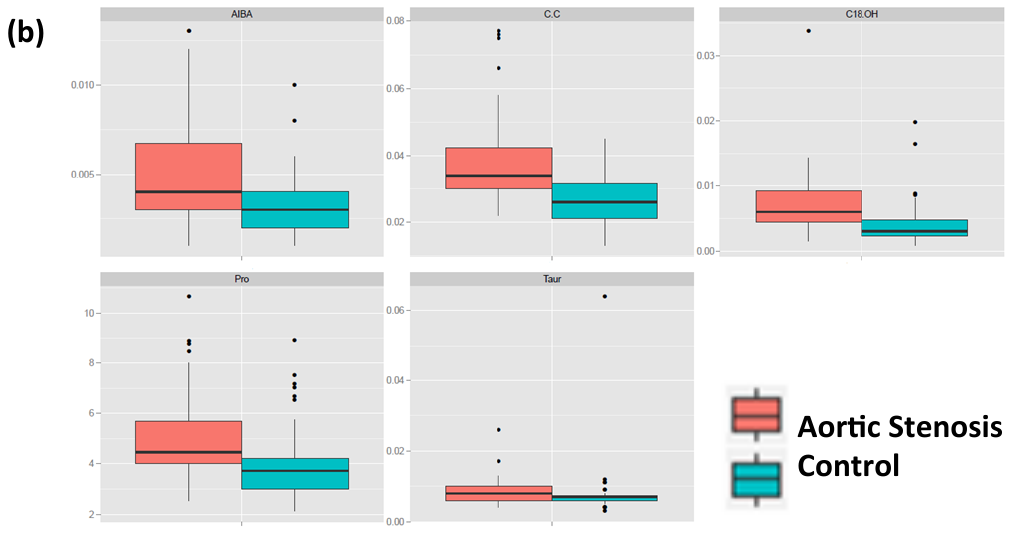

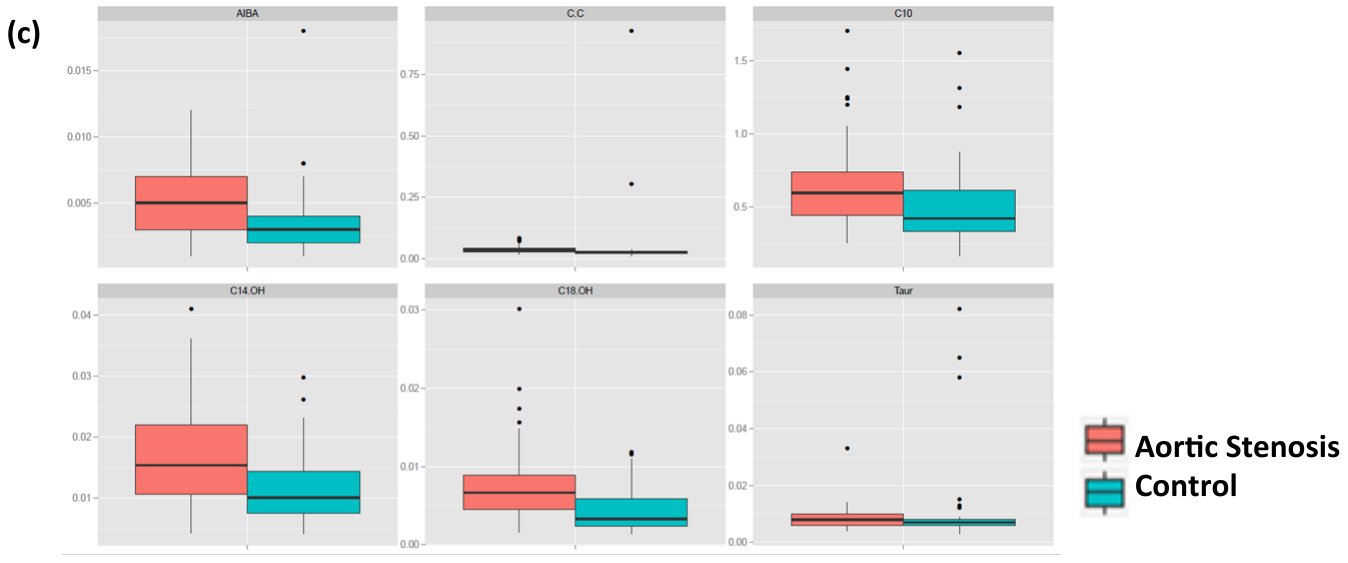

**Supplementary Figure 5 Metabolites** selected using random forest and their changes are shown in the box plot for control vs. hypertrophic cardiomyopathy: a) AR site b) CS site c) FV site**.**

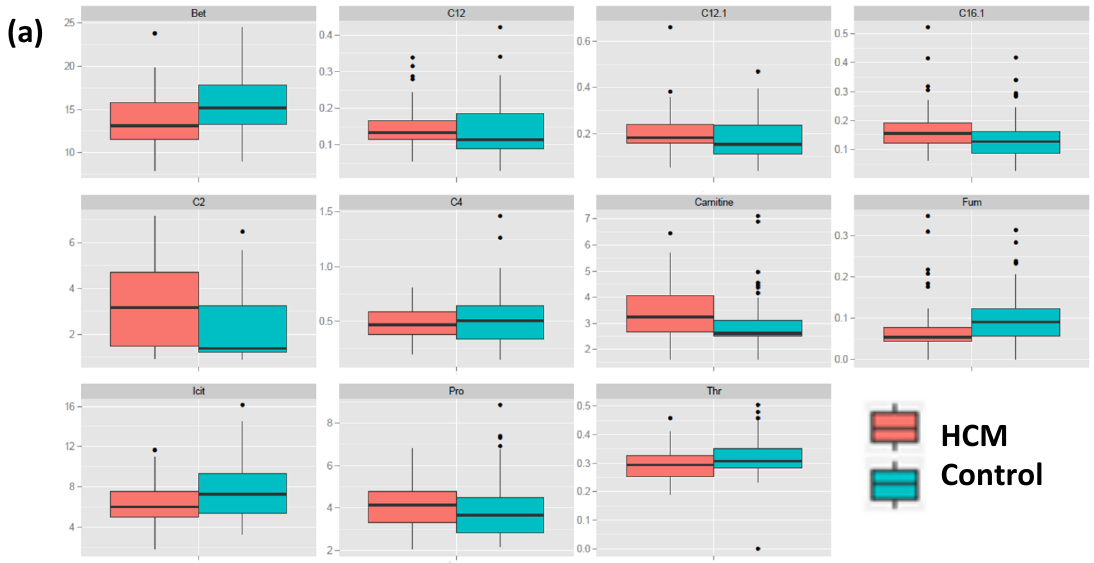

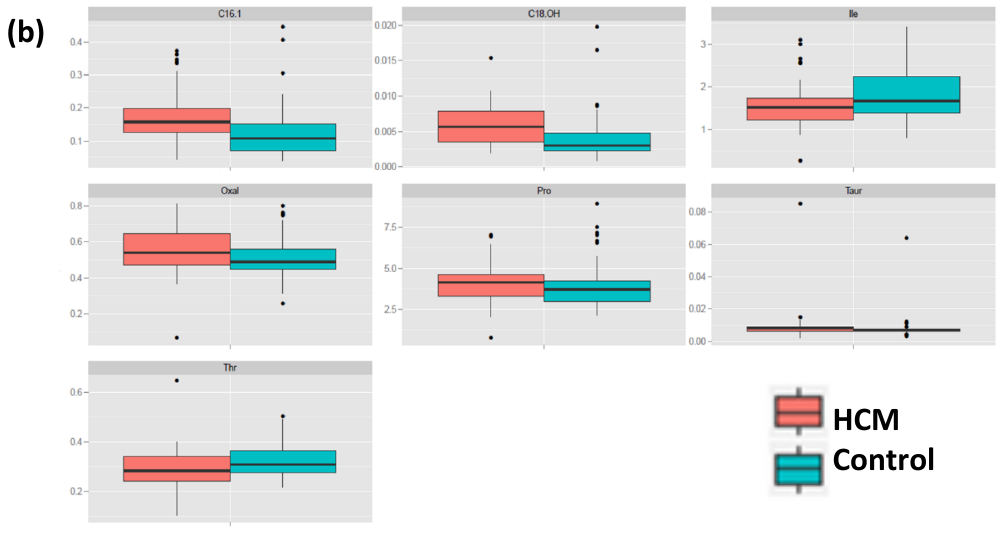

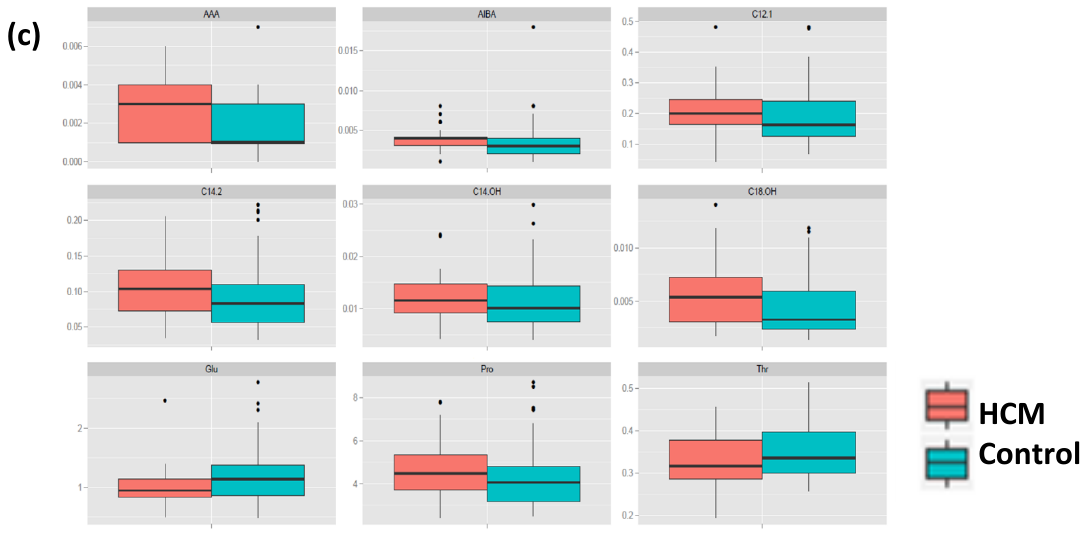

**Supplementary Figure 6** Metabolites selected using random forest and their changes are shown in the box plot for hypertrophic cardiomyopathy vs. aortic stenosis: a) AR site b) CS site c) FV site**.**

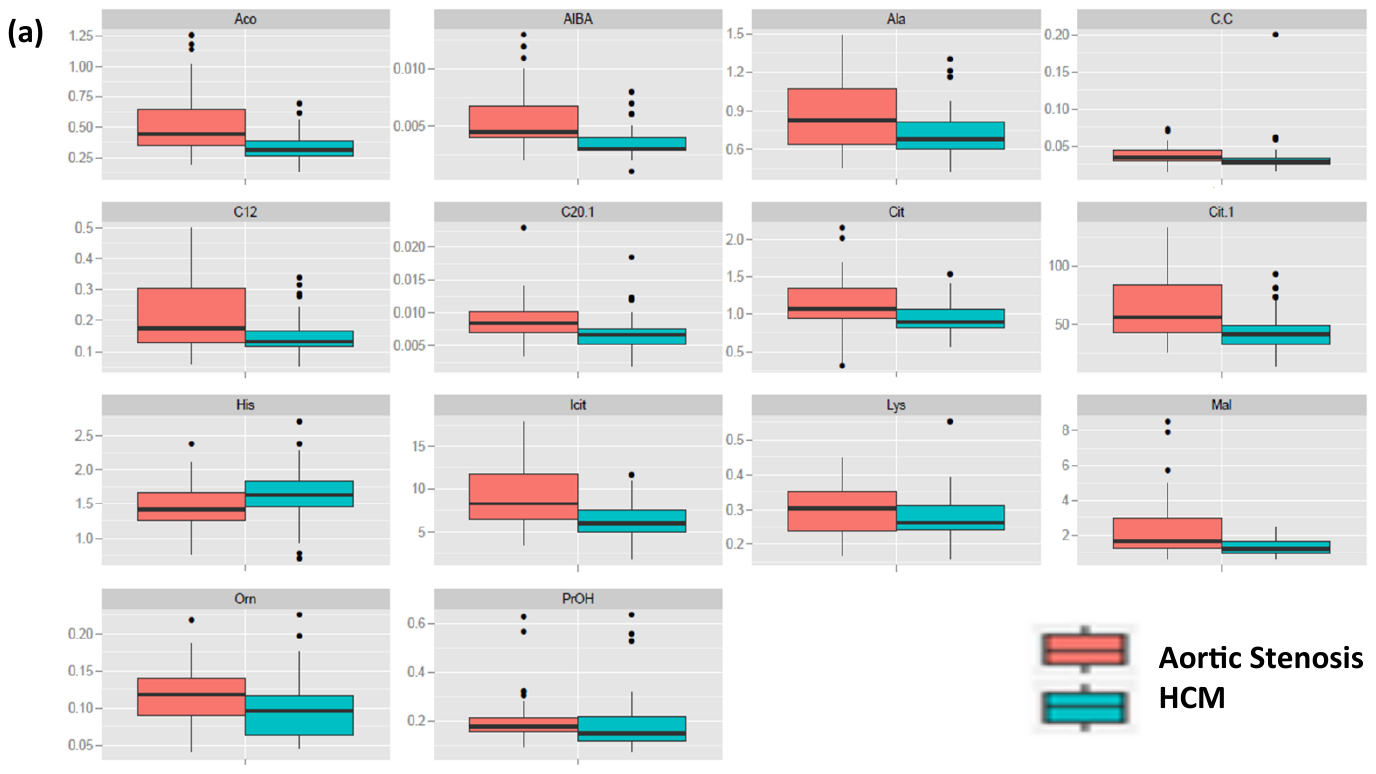

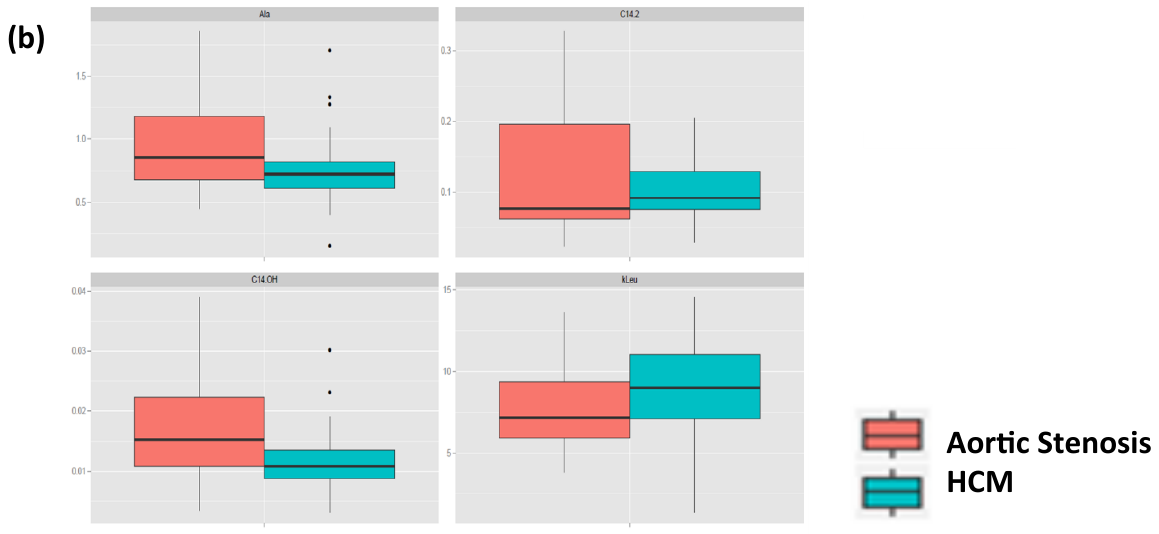

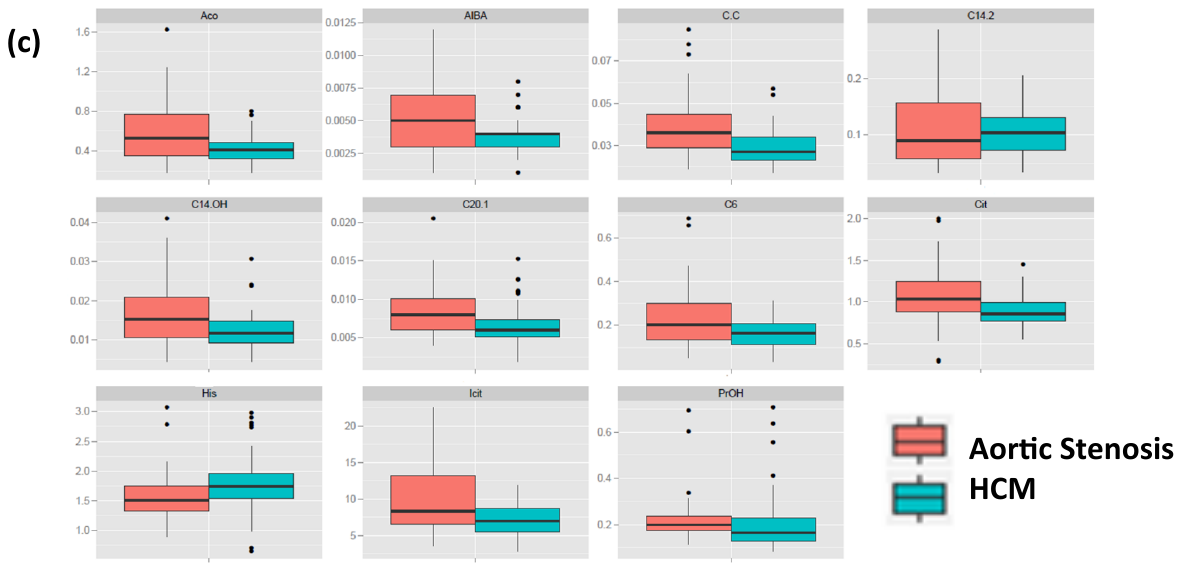

**Supplementary Figure 7** a) ROC values are plotted after metabolites selected using random forest (RF) and PLS-DA. Metabolites Selected by RF resulted in higher ROC values compared to PLS-DA selected metabolites. b) Comparing the number of metabolites selected in PLS-DA and random forest in different disease comparison: control vs aortic stenosis (AS), control vs hypertrophic cardiomyopathy (HCM) and AS vs HCM.

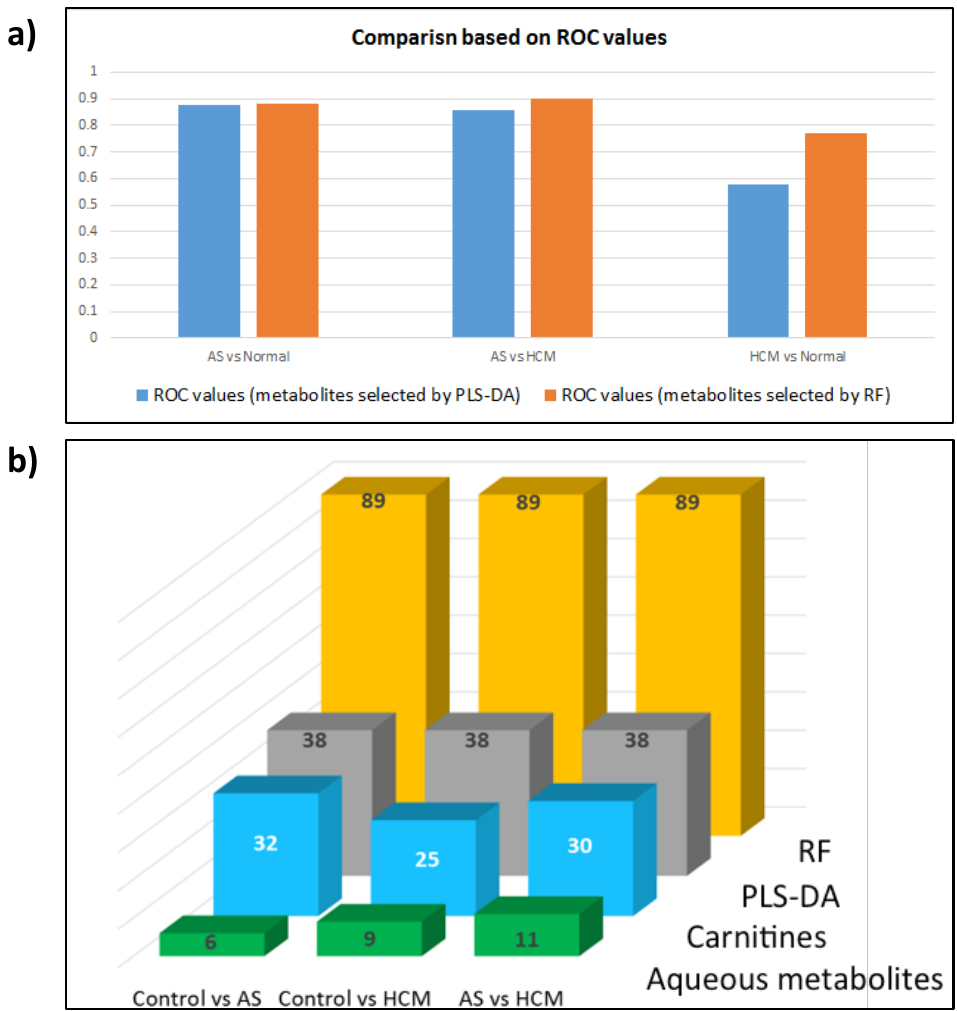

**Supplementary Figure 8** Stratifying models by age. To address the issue that the aortic stenosis group has a greater mean age than the other two groups we prepared models using only samples from those aged over 70. (**a**) Numbers of individuals in each classification according to age over 70, gender and disease status. (**b**) PLS-DA plot of aortic stenosis compared with controls for those over 70 (R^2^X=56.1%, R^2^Y=68.3%, Q^2^= 45.7%) (**c**) Random permutation test to validate the model in b. (**d**) In total 33 metabolites are selected and ranked based on VIP scores from PLS-DA model. Random forest selected less number of metabolites, which are shown in blue. Metabolites displayed in bold and italicised are common to analysis performed before and after age and gender stratification.

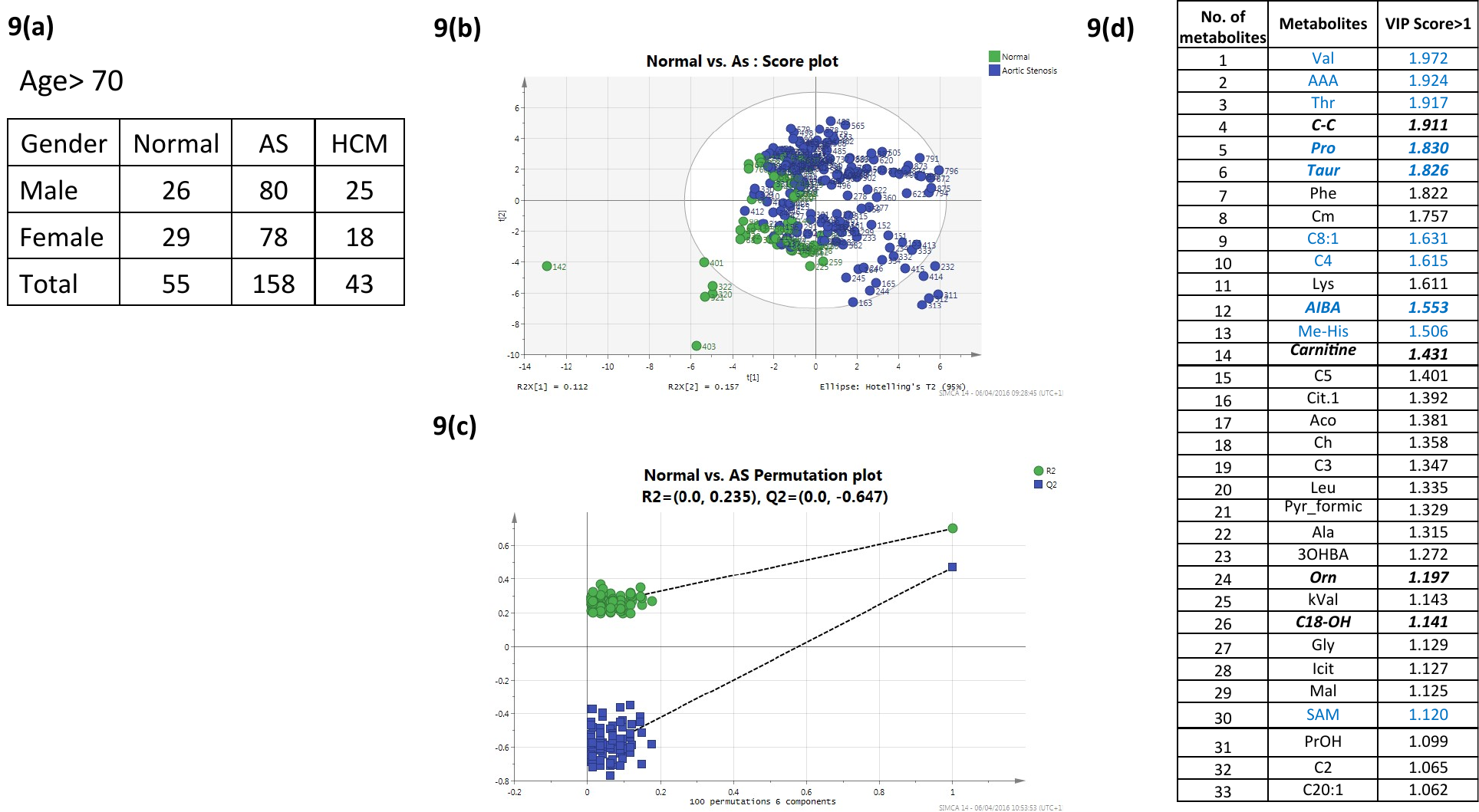

**Supplementary Figure 9** Selected metabolites identified by random forest models were used to generate ROC curves for each pairwise comparison and each site of blood sampling. Key: A: AUC > 0.9 (shown grey), B: AUC > 0.80 (shown red), AUC > 0.70 (shown green).

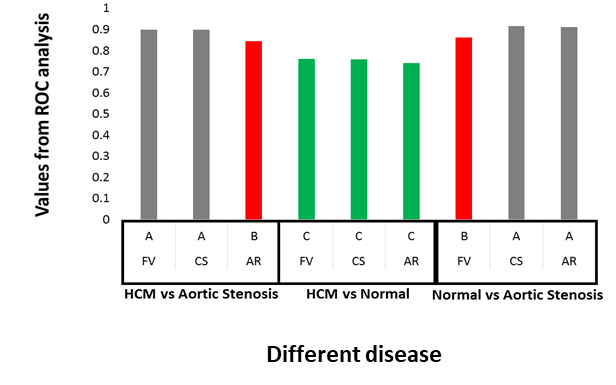

**Supplementary Figure 10** The heat map depicts the expression pattern of genes from the top three significantly enriched metabolic pathways found on pathway enrichment analysis between the aortic stenosis and control cohorts: a) Adipogenesis c) PPAR d) PPARα/RXRα activation. The expression data are centred on zero for each gene with red indicating a sample has higher than average expression for a given gene while green indicates lower than average expression.

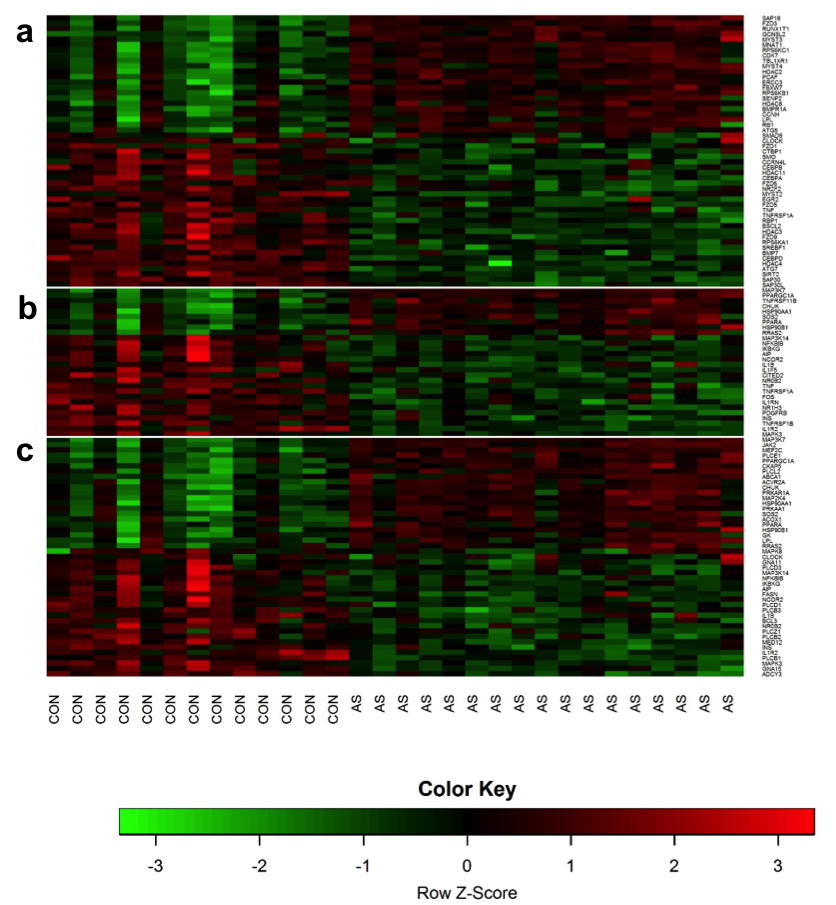

**Supplementary Table 1** Table details the biochemical characteristics of the controls, aortic stenosis and hypertrophic cardiomyopathy (HCM) cohorts. On a baseline biochemical analysis there were no difference in haemoglobin, total cholesterol and liver function tests between the 3 cohorts. The patients with aortic stenosis had a significantly lower eGFR (estimated glomerular filtration rate) and higher concentration of creatinine (89.8 µmol/L). Differences between the groups statistically analysed using ANOVA with Bonferroni’s post-test for multiple comparisons. Statistical significance was defined as a *P* < 0.05.

|  | | **Controls**  **(n = 38)** | | **Aortic stenosis**  **(n = 36)** | | | **HCM**  **(n = 35)** | | ***P* value** |
| --- | --- | --- | --- | --- | --- | --- | --- | --- | --- |
| Haemoglobin, g/dL | | 13.6 ± 1.8 | | 13.3 ± 1.5 | | | 13.9 ± 1.4 | | 0.15 |
| Creatinine, µmol/L | | 71.5 ± 13.3 | | 89.8 ± 31.8 | | | 84.6 ± 25.4 | | 0.008 |
| eGFR, ml/min/1.73m^2^ | | 100.8 ± 22.6 | | 72.2 ± 23.5 | | | 86.0 ± 23.0 | | <0.0001 |
| Cholesterol, mmol/L | | 4.3 ± 1.3 | | 4.2 ± 1.5 | | | 4.3 ± 1.1 | | 0.6 |
| Bilirubin, μmol/L | | 10 (8 - 13) | | 10 (8 - 18) | | | 11 (8 - 16) | | 0.6 |
| ALT, IU/L | | 25 (18 - 35) | | 18 (13 - 21) | | | 21 (14 - 31) | | 0.005 |
| ALP, IU/L | | 165 (133 - 187) | | 149 (131 - 225) | | | 166 (127 - 230) | | 0.9 |
| Albumin, g/L | | 44 (41 - 47) | | 43 (41 - 45) | | | 44 (42 - 45) | | 0.2 |

*Values are mean ± SD or median (quartiles 1 to 3). ALT = alanine transaminase. ALT = alanine transaminase; ALP = alkaline phosphatase; Cholesterol = total cholesterol.*

**Supplementary Table 2** Table details the baseline hemodynamic and echocardiographic characteristics of control, aortic stenosis and hypertrophic cardiomyopathy (HCM) patients. Patients in all three groups had similar baseline heart rate and systolic blood pressure measurements (supplementary table 2). Echocardiography showed that although the left ventricular (LV) ejection fraction was normal in all the patients, it was significantly higher in patients with hypertrophic cardiomyopathy (HCM) when compared with aortic stenosis (AS) (*P* = 0.005). Septal hypertrophy was most marked in patients with HCM (1.8 ± 0.4 cm), having an eccentric hypertrophy pattern as compared to patients with AS who had concentric hypertrophy and the highest LV mass index. Diastolic function was impaired in patients with aortic stenosis with a low e’ (6.5 cm/s) and a raised E/e’ ratio = 14.7. These patients had severe AS with a mean AS jet velocity of 419 cm/s (meters per second), aortic valve area of 0.7 cm^2^ and an indexed valve area of 0.4 cm^2^/m^2^. Differences between the groups analysed statistically using ANOVA with Bonferroni’s post-test for multiple comparisons. Statistical significance was defined as a *P* < 0.05.

|  |  | **Controls**  **(n = 22)** | **Aortic stenosis**  **(n = 34)** | **HCM**  **(n = 31)** | ***P* value** |
| --- | --- | --- | --- | --- | --- |
| Heart rate, beats/min |  | 68 (60 - 79) | 63 (57 - 74) | 64 (58 - 70) | 0.09 |
| Systolic BP, mmHg |  | 128 ± 21 | 134 ± 30 | 129 ± 22 | 0.6 |
| Diastolic BP, mmHg |  | 70 ± 12 | 63 ± 9 | 78 ± 14 | <0.0001 |
| LVEF (%) |  | 61 ± 10 | 56 ±12 | 66 ± 11 | 0.005 |
| LVIDd, mm |  | 4.6 ± 0.5 | 4.8 ± 0.9 | 4.3 ± 0.7 | 0.04 |
| IVSd, mm |  | 1.2 ± 0.2 | 1.4 ± 0.2 | 1.8 ± 0.4 | <0.0001 |
| PWd, mm |  | 1.1 ± 0.2 | 1.4 ± 0.2 | 1.2 ± 0.3 | 0.0001 |
| LVIDs, mm |  | 2.9 ± 0.5 | 3.2 ± 1.0 | 2.6 ± 0.6 | 0.01 |
| LV mass index, g/m^2^ |  | 96 ± 20 | 149 ± 50 | 132 ± 35 | <0.0001 |
| E/A ratio |  | 0.8 (0.7 - 1.1) | 0.8 (0.7 - 1.8) | 0.9 (0.7 - 1.4) | 0.6 |
| DT, m/s |  | 224 ± 53 | 230 ± 87 | 209 ± 61 | 0.7 |
| e’, cm/s |  | 8.1 ± 2.1 | 6.5 ± 1.5 | 6.2 ± 1.7 | 0.009 |
| E/e’ ratio |  | 8.2 ± 1.9 | 14.7 ± 6.6 | 10.8 ± 3.6 | 0.01 |
| AS jet velocity, cm/s |  | 127 (122 - 154) | 419 (382 - 485) | 136 (123 - 197) | <0.0001 |
| AVA, cm^2^ |  |  | 0.7 ± 0.2 |  |  |
| Indexed AVA, cm^2^/m^2^ |  |  | 0.4 ± 0.1 |  |  |

*Values are mean ± SD, percentages or median (quartiles 1 to 3). LV = left ventricular; LA = left atrial; LVEF = left ventricular ejection fraction; LVIDd = left ventricular internal diameter at end-diastole; IVSd = interventricular septum diameter (cm); PWd = posterior wall diameter;* *LVIDs = left ventricular internal diameter at end-systole; E = peak early diastolic mitral flow velocity; A = late diastolic mitral flow velocity;* *DT = E wave deceleration time; e’ = peak early diastolic mitral annular velocity (mean of septal and lateral); AS = aortic stenosis; AVA = aortic valve area.*

**Supplementary Table 3** Table details the trans-cardiac gradients for measured metabolites at baseline. To assess the uptake and export of metabolites by the heart the trans-cardiac gradients were calculated by subtracting the metabolite concentrations measured in blood plasma from the aortic root from that detected in the coronary sinus. Values are mean ± standard error of the mean. Key: * *P <* 0.05, ** *P* <0.01, *** *P <* 0.001 for significant difference from zero across the gradient with positive values indicative of higher concentrations at the aortic root. To compare between cohorts an ANOVA test was performed followed by a Welch post-test. # *P* <0.05, ## *P <* 0.01, ### *P <* 0.001 for a significant difference between aortic stenosis and control, ^ *P* < 0.05 for significant difference between hypertrophic cardiomyopathy and control, & *P* < 0.05 for significant difference between hypertrophic cardiomyopathy and aortic stenosis.

|  | **Controls**  **(n = 38)** | **Aortic stenosis**  **(n = 36)** | **HCM**  **(n = 35)** | ***P* value** |
| --- | --- | --- | --- | --- |
| *Lysine* | -0.0061 ± 0.0068 | **-0.013 ± 0.006*** | -0.048 ± 0.052 |  |
| *Ornithine* | -0.0028 ± 0.0031 | **-0.010 ± 0.005*** | -0.035 ± 0.037 |  |
| *Glutamate* | **1.019 ± 0.134*** | **0.653 ± 0.086*** | **0.661 ± 0.169*** | # |
| *Pyruvate* | **0.0783 ± 0.0224*** | **0.115 ± 0.032**** | 0.0063 ± 0.019 | ^, && |
| *Malate* | -0.062 ± 0.075 | 0.108 ± 0.142 | -0.249 ± 0.084 | & |
| *Isocitrate* | 0.166 ± 0.331 | 0.292 ± 0.399 | **-1.081 ± 0.383*** | ^, & |
| *Methionine* | 4.204 ± 2.421 | 2.274 ± 3.291 | -4.250 ± 3.144 | ^ |
| *Succinate* | **-0.181 ± 0.033***** | **-0.119 ± 0.038*** | **-0.146 ± 0.037**** |  |
| *Aconitate* | 0.0094 ± 0.0278 | 0.0052 ± 0.0344 | **-0.0584 ± 0.0213*** |  |
| *3-hydroxy butyric acid* | **-0.055± 0.027*** | **-0.0655 ± 0.0320*** | **-0.119 ± 0.030**** |  |
| *2-hydroxy butyric acid* | **0.578 ± 0.182**** | 0.497 ± 0.163 | -0.020 ± 0.151 | & |
| *Leucine* | **0.114 ± 0.042*** | 0.064 ± 0.044 | 0.071 ± 0.044 |  |
| *α-ketoisovalerate* | **0.727 ± 0.135***** | **0.781 ± 0.142***** | 0.282 ± 0.118 | ^, && |
| *Keto-isoleucine (KIso)* | **1.350 ± 0.248***** | **1.121 ± 0.316**** | 0.245 ± 0.332 | ^ |
| *Keto-leucine (KLeu)* | **1.766 ± 0.427***** | **2.059 ± 0.535**** | 0.249 ± 0.390 | ^^^, &&& |
| *Carnitine* | 0.0297 ± 0.050 | -0.476 ± 0.194 | 0.063 ± 0.173 | #, & |
| *C4-carnitine* | **-0.0431 ± 0.0172*** | 0.028 ± 0.040 | -0.0406 ± 0.0300 |  |
| *C8-carnitine* | -0.00058 ± 0.0154 | -0.021 ± 0.021 | 0.00688 ± 0.0150 | # |
| *C10-carnitine* | 0.0176 ± 0.027 | **-0.082 ± 0.029*** | 0.0101 ± 0.025 | #, & |
| *C12:1-carnitine* | 0.00085 ± 0.0105 | **-0.020 ± 0.0094*** | 0.00043 ± 0.0111 |  |
| *C14-OH-carnitine* | -0.00034 ± 0.00066 | -0.001 ± 0.00062 | 0.0014 ± 0.00090 | & |
| *C16:1-OH-carnitine* | 0.00025 ± 0.00036 | **0.00099 ± 0.00038*** | 0.00070 ± 0.00055 |  |
| *C20:2-carnitine* | **0.00009 ± 0.00024****  ****** | 0.00011 ± 0.00015 | 0.00016 ± 0.00028 |  |
| *C20:1-carnitine* | 0.00057 ± 0.00036 | 0.00062 ± 0.00038 | **0.00093 ± 0.00040***  ***** |  |

*Values are mean ± SD.*

**Supplementary Table 4** Table details the trans-cardiac gradients for metabolites measured during pacing. To assess the uptake and export of metabolites by the heart the trans-cardiac gradients were calculated by subtracting the metabolite concentrations measured in blood plasma from the aortic root from that detected in the coronary sinus. Values are mean ± standard error of the mean. Key: * *P* < 0.05, ** *P* < 0.01, *** *P* < 0.001 for significant difference from zero across the gradient with positive values indicative of higher concentrations at the aortic root. To compare between groups an ANOVA test was performed followed by a Welch post-test. # *P* < 0.05, ## *P* < 0.01, ### *P* < 0.001 for a significant difference between aortic stenosis and control, ^ *P* < 0.05 for significant difference between hypertrophic cardiomyopathy and control, & *P* < 0.05 for significant difference between hypertrophic cardiomyopathy and aortic stenosis.

| **Metabolite** | **Controls** | **Aortic stenosis** | **HCM** | ***P* value** |
| --- | --- | --- | --- | --- |
| *Histidine* | 0.113 ± 0.114 | 0.126 ± 0.064 | **0.0144 ± 0.063 *** |  |
| *Hydroxy-proline* | 0.00047 ± 0.0110 | **0.0157 ± 0.0076 *** | 0.0114 ± 0.0068 |  |
| *Glutamate* | **0.566 ± 0.202 *** | **0.424 ± 0.102 **** | **0.562 ± 0.133 **** |  |
| *Betaine* | **1.062 ± 0.497 *** | 0.195 ± 0.357 | 0.455 ± 0.417 |  |
| *Pyruvate* | **0.0810 ± 0.0323 *** | **0.0822 ± 0.0344 *** | 0.0117 ± 0.0349 |  |
| *Valine* | 0.178 ± 0.105 | **0.114 ± 0.0507 *** | 0.142 ± 0.0957 |  |
| Cytidine *(Cyd)* | 0.00100 ± 0.00072 | -0.00057 ± 0.00044 | -0.00066 ± 0.00048 | ^ |
| *Succinate* | **-0.161 ± 0.052 **** | **-0.169 ± 0.046 ***** | **-0.181 ± 0.062 **** |  |
| *2-hydroxy butyric acid* | **0.630 ± 0.261 *** | **0.341 ± 0.144 *** | **0.463 ± 0.214 *** |  |
| *Leucine* | **0.183 ± 0.083 *** | **0.112 ± 0.040 *** | 0.156 ± 0.081 |  |
| *Methyl-cytosine* | 0.0762 ± 0.0779 | 0.00009 ± 0.00005 | **0.0156 ± 0.00005 **** |  |
| *Uridine* | 0.0451 ± 0.0453 | **-0.00083 ± 0.00031 *** | -0.00144 ± 0.00074 |  |
| *α-ketoisovalerate* | **0.817 ± 0.197 ***** | **0.644 ± 0.165 ***** | **0.629 ± 0.168 **** |  |
| *Keto-isoleucine (KIso)* | **1.131 ± 0.507 *** | **0.701 ± 0.325 *** | **0.887 ± 0.360 *** |  |
| *Keto-leucine (KLeu)* | **2.324 ± 0.601 ***** | **1.532 ± 0.434 **** | **1.565 ± 0.587 *** |  |
| *Carnitine* | **0.0234 ± 0.104 *** | 0.127 ± 0.129 | 0.102 ± 0.117 |  |
| *C8-carnitine* | -0.00384 ± 0.0192 | -0.0614 ± 0.0140 | -0.0172 ± 0.0096 | & |
| *C4-DC-carnitine* | **0.00112 ± 0.0005 *** | 0.00044 ± 0.00074 | 0.00031 ± 0.00048 |  |
| *C12:1-carnitine* | 0.00053 ± 0.0095 | **-0.0201 ± 0.0094** | -0.00469 ± 0.0090 |  |
| *C8-DC-carnitine* | 0.00465 ± 0.00370 | **0.0071 ± 0.0030 *** | 0.00012 ± 0.0013 |  |
| *C14-OH-carnitine* | **0.00117 ± 0.00052 *** | **0.00202 ± 0.00096 *** | 0.0014 ± 0.00084 |  |
| *C18:2-OH-carnitine* | 0.00075 ± 0.00051 | -0.00081 ± 0.00058 | -0.00094 ± 0.00057 | #, ^ |

*Values are mean ± SD.*

**Supplementary Table 5** Table details the clinical and demographic characteristics of the controls and aortic stenosis cohorts in the validation group. Differences between the groups statistically analysed using Student’s t-test. Statistical significance was defined as a *P* < 0.05.

|  | **Controls**  **(n = 23)** | **Aortic**  **Stenosis**  **(n = 27)** | **HCM**  **(n = 30)** | ***P* value** |
| --- | --- | --- | --- | --- |
| Age (years) | 61 (41- 65) | 71 (65 - 76) | 53 (41 - 58) | <0.0001 |
| Men, n (%) | 61 | 76 | 67 | 0.52 |
| LVEF (%) | 69 ± 11 | 74 ± 6 | 77 ± 8 | 0.003 |

*Values are mean ± SD, percentages or median (quartiles 1 to 3). LVEF = left ventricular (LV) ejection fraction.*

**Supplementary Table 6** Table details the demographic characteristics of the controls and aortic stenosis patients selected for transcriptomic analysis. While AS patients undergoing microarray analysis were significantly younger than those in the discovery metabolomics cohort (68 vs 81 years, *P* < 0.0001), their mean age was comparable to those in the validation cohort (68 vs 71 years, *P* = 0.45). The mean age of control subjects recruited for microarray analysis was comparable to those in the discovery cohort (71 vs 62, *P* = 0.20), but were significantly older than the validation cohort (71 vs 61, *P* < 0.004). There were no differences in gender distribution between the cohorts. Differences between the groups statistically analysed using either two-tailed Students t-test, Mann-Whitney U-test or Chi-square test. *P* < 0.05 was considered significant.

|  | **Controls**  **(n = 13)** | **Aortic**  **Stenosis**  **(n = 17)** | ***P* value** |
| --- | --- | --- | --- |
| Age (years) | 71 (61 - 73) | 68 (64 - 72) | 0.84 |
| Men, n (%) | 77 | 71 | 1.0 |

*Values are mean ± SD, percentages or median (quartiles 1 to 3).*

**Supplementary Table 7** Table details the list of Ingenuity Canonical pathways in an order of decreasing significance on enrichment analysis between the aortic stenosis and control cohorts. This table is attached as a separate excel file.

**Supplementary Table 8** Table details the differential expression of selected genes, which regulate the metabolomic pathways, between the aortic stenosis and control cohorts; the genes are listed in an order of decreasing significance,. Differences between the groups statistically analysed using either two-tailed Student’s t-test or Mann-Whitney U-test. Statistical significance was defined as an adjusted *P* < 0.05.

| **Gene** | **Log2-fold change** | **Average Expression** | **t** | **Adjusted *P* value** |
| --- | --- | --- | --- | --- |
| *PCK2* | -0.94304 | 4.12738 | -7.07753 | 2.06E-06 |
| *ACOT4* | -0.66283 | 6.31591 | -7.01586 | 2.34E-06 |
| *CYP27A1* | -0.82034 | 8.36832 | -6.59802 | 5.53E-06 |
| *PFKFB2* | 1.04581 | 8.96158 | 6.44123 | 7.62E-06 |
| *PPARGC1A* | 0.88620 | 10.73207 | 6.40810 | 8.20E-06 |
| *ADCY3* | -0.70386 | 9.56585 | -6.22089 | 1.20E-05 |
| *ACOT7* | -0.67240 | 6.30104 | -5.77249 | 3.08E-05 |
| *ACADSB* | 1.15181 | 6.49135 | 5.67472 | 3.77E-05 |
| *ACADM* | 1.22082 | 11.41422 | 5.64463 | 4.04E-05 |
| *INS* | -0.75475 | 4.11575 | -5.58992 | 4.50E-05 |
| *ACAD8* | -0.89748 | 3.77865 | -5.57073 | 4.68E-05 |
| *ACSL3* | 1.03796 | 6.79234 | 5.45688 | 6.00E-05 |
| *GPAM* | 0.86156 | 7.52223 | 5.37754 | 7.11E-05 |
| *HSD17B10* | -0.44841 | 10.44612 | -5.33970 | 7.71E-05 |
| *ACAD11* | 0.85404 | 9.44550 | 5.27139 | 9.03E-05 |
| *SLC27A5* | -0.61087 | 6.17650 | -5.15540 | 0.00012 |
| *ABCD3* | 0.83774 | 7.28104 | 5.12946 | 0.00012 |
| *DGAT2* | -0.58644 | 5.05769 | -4.95904 | 0.00019 |
| *FABP6* | -1.03060 | 5.58404 | -4.88296 | 0.00022 |
| *CROT* | 0.78562 | 6.53046 | 4.76083 | 0.00029 |
| *G6PC3* | -0.54337 | 10.06723 | -4.70685 | 0.00033 |
| *PDK2* | -0.56290 | 8.45308 | -4.62577 | 0.00040 |
| *ACOT8* | -0.60991 | 7.23119 | -4.61819 | 0.00040 |
| *ELOVL1* | -0.42448 | 7.12705 | -4.61698 | 0.00040 |
| *PRKAA1* | 0.58254 | 7.13844 | 4.61377 | 0.00041 |
| *HSD17B4* | 0.80000 | 10.55513 | 4.58885 | 0.00043 |
| *OAZ3* | -0.53083 | 4.63217 | -4.38173 | 0.00069 |
| *ACOX1* | 0.50306 | 7.11018 | 4.29867 | 0.00084 |
| *PPARA* | 0.43225 | 5.25548 | 4.23713 | 0.00097 |
| *ETFDH* | 0.70069 | 11.36640 | 4.07806 | 0.00139 |
| *GK* | 0.73248 | 5.92392 | 3.86974 | 0.00224 |
| *FASN* | -0.55726 | 8.56746 | -3.82025 | 0.00250 |
| *AIP* | -0.44586 | 9.41173 | -3.80994 | 0.00256 |
| *LPL* | 0.65686 | 12.19096 | 3.80696 | 0.00258 |
| *DHRS3* | -0.35571 | 9.76141 | -3.68391 | 0.00341 |
| *AGPAT7* | -0.56337 | 6.14112 | -3.56367 | 0.00445 |
| *HK2* | 0.91035 | 6.26182 | 3.55945 | 0.00449 |
| *ASS1* | -0.55970 | 7.75036 | -3.53246 | 0.00478 |
| *CPT2* | -0.36771 | 9.53403 | -3.51489 | 0.00496 |
| *ETFB* | -0.32112 | 11.98809 | -3.48563 | 0.00529 |
| *PDK1* | -0.46505 | 3.84936 | -3.44692 | 0.00577 |
| *PRKAG2* | 0.42774 | 9.49158 | 3.41002 | 0.00625 |
| *CYP4X1* | -0.38248 | 5.49362 | -3.40454 | 0.00632 |
| *ECH1* | -0.37216 | 12.99107 | -3.28711 | 0.00820 |
| *SLC25A11* | -0.40718 | 10.26882 | -3.26380 | 0.00863 |
| *CPT1A* | -0.32446 | 5.45122 | -3.21326 | 0.00962 |
| *DECR2* | -0.38335 | 8.08883 | -3.16932 | 0.01061 |
| *CD36* | 0.55447 | 11.98284 | 3.16422 | 0.01071 |
| *ELOVL6* | 0.44058 | 4.22791 | 3.13301 | 0.01148 |
| *HMGCS2* | -2.13096 | 7.84109 | -3.13027 | 0.01153 |
| *PRKAG1* | -1.88024 | 4.18373 | -3.11811 | 0.01183 |
| *CYP4B1* | -1.22582 | 6.05511 | -3.04842 | 0.01371 |
| *PRKAB1* | -0.32354 | 7.32820 | -3.03143 | 0.01417 |
| *CRAT* | -0.33899 | 9.40907 | -3.01953 | 0.01455 |
| *ACO2* | -0.28708 | 11.15327 | -2.99506 | 0.01533 |
| *ADH1B* | -0.70120 | 6.37155 | -2.93448 | 0.01743 |
| *ADH1C* | -0.68321 | 5.86069 | -2.81868 | 0.02218 |
| *EHHADH* | 0.36969 | 5.88289 | 2.73338 | 0.02636 |
| *SLC25A10* | -0.34104 | 7.12889 | -2.60199 | 0.03422 |
| *ACOT1* | -0.44481 | 10.34146 | -2.52736 | 0.03952 |
| *AGPAT5* | -0.51137 | 6.01440 | -2.42389 | 0.04820 |
| *DGAT1* | -0.32047 | 7.30025 | -2.38073 | 0.05233 |
| *ACADS* | -0.37541 | 8.85812 | -2.36254 | 0.05426 |
| *SLC25A20* | -0.28770 | 9.17171 | -2.35945 | 0.05455 |
| *CPT1C* | -0.32165 | 5.96773 | -2.26426 | 0.06530 |
| *PNPLA2* | -0.26184 | 9.29186 | -1.89113 | 0.12584 |
| *ARG2* | -0.25157 | 4.82596 | -1.65742 | 0.18194 |
| *LPIN2* | 0.22351 | 5.92380 | 1.64474 | 0.18557 |
| *CPT1B* | -0.21679 | 10.03904 | -1.59220 | 0.20107 |
| *VNN1* | -0.30393 | 3.73222 | -1.41625 | 0.25971 |
| *CLOCK* | -0.19151 | 7.08781 | -1.30471 | 0.30195 |
| *SLC2A4* | -0.12689 | 6.33656 | -1.20548 | 0.34168 |
| *SLC2A4* | -0.12689 | 6.33656 | -1.20548 | 0.34168 |
| *PRKAA2* | 0.16165 | 6.15386 | 1.13446 | 0.37257 |
| *PDK4* | -0.08426 | 10.12328 | -0.19316 | 0.89302 |
| *ELOVL2* | 0.01364 | 4.67620 | 0.08352 | 0.95446 |

**Supplementary Table 9** Table details the top 25 genes, which are most significantly different between the aortic stenosis and control cohorts. Differences between the groups statistically analysed using either two-tailed Student’s t-test or Mann-Whitney U-test. Statistical significance was defined as an adjusted *P* < 0.05.

| **Gene** | **Log2-fold change** | **Average Expression** | **t** | **Adjusted *P* value** |
| --- | --- | --- | --- | --- |
| *FTHL8* | -1.53336 | 8.33470 | -15.02494 | 4.16E-12 |
| *FTHL11* | -1.26639 | 8.25951 | -13.52830 | 4.24E-11 |
| *LOC388344* | -2.66720 | 4.49422 | -13.25579 | 5.02E-11 |
| *FTHL12* | -1.44102 | 9.16762 | -12.90101 | 8.05E-11 |
| *RPLP1* | -1.84308 | 8.23730 | -12.53364 | 1.44E-10 |
| *HS.554507* | 2.05492 | 6.04981 | 12.06465 | 3.41E-10 |
| *PTMA* | -1.83840 | 5.29332 | -11.95229 | 3.77E-10 |
| *LOC645138* | -1.46990 | 10.89377 | -11.83291 | 4.33E-10 |
| *LOC136143* | -1.27612 | 6.33176 | -11.68032 | 5.47E-10 |
| *METTL7A* | -2.00076 | 4.38889 | -11.52634 | 7.03E-10 |
| *LOC647100* | -1.36339 | 7.37649 | -11.41248 | 8.33E-10 |
| *RPL23* | -1.46369 | 9.86886 | -11.35477 | 8.74E-10 |
| *XAF1* | 1.31955 | 9.00462 | 11.21505 | 1.12E-09 |
| *SUMO2* | -1.31949 | 8.10139 | -10.88352 | 2.16E-09 |
| *ZFP36L1* | -1.52098 | 6.55967 | -10.85306 | 2.16E-09 |
| *LOC390354* | -1.55321 | 10.31584 | -10.85306 | 2.16E-09 |
| *RPL27A* | -1.59559 | 5.85772 | -10.78264 | 2.35E-09 |
| *LOC648622* | -1.29374 | 10.88555 | -10.77020 | 2.35E-09 |
| *FTHL2* | -1.03847 | 9.94768 | -10.74274 | 2.38E-09 |
| *LOC388076* | -1.02237 | 9.66789 | -10.56349 | 3.49E-09 |
| *NDUFB9* | -1.17064 | 12.36196 | -10.26455 | 6.75E-09 |
| *LSM6* | -0.99919 | 4.35139 | -10.25811 | 6.75E-09 |
| *C1ORF212* | -1.11621 | 5.61148 | -10.08422 | 9.96E-09 |
| *LOC644039* | -1.00891 | 13.51888 | -10.06261 | 1.01E-08 |
| *FTHL8* | -1.53336 | 8.33470 | -15.02494 | 4.16E-12 |
| *FTHL11* | -1.26639 | 8.25951 | -13.52830 | 4.24E-11 |
| *LOC388344* | -2.66720 | 4.49422 | -13.25579 | 5.02E-11 |
| *FTHL12* | -1.44102 | 9.16762 | -12.90101 | 8.05E-11 |
